# Untangling intelligence, psychopathy, antisocial personality disorder, & conduct problems: A meta-analytic review

**DOI:** 10.1101/100693

**Authors:** Olga Sánchez de Ribera, Nicholas Kavish, Ian M. Katz, Brian B. Boutwell

## Abstract

Substantial research has investigated the association between intelligence and psychopathic traits. The findings to date have been inconsistent and have not always considered the multi-dimensional nature of psychopathic traits. Moreover, there has been a tendency to confuse psychopathy with other closely related, clinically significant disorders. The current study represents a meta-analysis conducted to evaluate the direction and magnitude of the association of intelligence with global psychopathy, as well as its factors and facets, and related disorders (Antisocial Personality Disorder, Conduct Disorder, and Oppositional Defiant Disorder). Our analyses revealed a small, significant, negative relationship between intelligence and total psychopathy (*r* = -.07, *p* = .001). Analysis of factors and facets found differential associations, including both significant positive (e.g., interpersonal facet) and negative (e.g., affective facet) associations, further affirming that psychopathy is a multi-dimensional construct. Additionally, intelligence was negatively associated with Antisocial Personality Disorder (*r* = -.13, *p* = .001) and Conduct Disorder (*r* = -.11, *p* = .001), but positively with Oppositional Defiant Disorder (*r* = .06, *p* = .001). There was significant heterogeneity across studies for most effects, but the results of moderator analyses were inconsistent. Finally, bias analyses did not find significant evidence for publication bias or outsized effects of outliers.

---

Psychopathy and intelligence represent two psychological constructs that have been studied extensively over the last several decades. Large bodies of psychometric work have consistently supported the reliability and validity of both concepts (Carrol, 1993; Hare et al., 1990; Kranzler & Jensen, 1991; Salekin, Rogers, & Sewell, 1996). General intelligence is one of the most studied traits in all of psychology and has nearly a century of research related to its measurement, development, and etiological underpinnings (Gottfredson, 2002; Ritchie, 2015). Psychopathy, while representing a more recently defined psychological construct (Cleckley, 1941), is nonetheless psychometrically robust, and research continues to shed light on its etiology and development across the life course.

Of particular interest to the current study, however, is a more recent line of research examining the association between indicators of intelligence and psychopathic traits. The last decade, in fact, has seen a sharp increase in studies examining the association between general intelligence and psychopathy, with some evidence suggesting that lower intelligence scores are correlated with increased psychopathic tendencies (e.g., DeLisi, Vaughn, Beaver, & Wright, 2010; Vitacco, Neumann, & Wodeshuk, 2008). To date, however, the results gleaned from this growing body of research have been somewhat mixed, with some studies, such as those cited above, finding evidence of a negative relationship between the two variables, and other studies failing to find such an effect.

The primary goal of the current study is to systematically review the literature in order to better understand the pattern of findings to date. To the extent that psychopathy covaries with intelligence (regardless of the direction of the association), it may provide insight into the development of both outcomes. Specifically, if intelligence and psychopathy are developmentally or clinically associated, future research could attempt to explore whether they are causally related in any manner or further clarify the shared mechanisms underlying the emergence of both constructs. An understanding of their (potentially) shared etiology, moreover, could have implications for intervention and perhaps even prevention.

Before progressing further, though, it is worth pointing out that while the primary goal of this study was to examine the association between intelligence and psychopathic traits, we also examine the association between intelligence and three other closely related antisocial constructs (antisocial personality disorder, conduct disorder, and oppositional defiant disorder). We do so, because these constructs are highly overlapping, yet they are not isomorphic. Thus, understanding their shared and unique associations with other important constructs (intelligence in this case), may further clarify the manner in which these disorders are both related and distinct from each other. Our rationale for including these constructs is further elaborated on in later portions of the introduction. First, though, we move to a more detailed overview of psychopathy as a clinical construct.

## Psychopathy

Unlike most clinical disorders that are characterized by a set of symptoms, psychopathy is commonly described as a cluster of *relatively* stable personality traits (Cleckley, 1941; Hare & Vertommen, 1991). The traits most often associated with psychopathy are callousness, remorselessness, lack of empathy, grandiosity, impulsivity, deceitfulness, and manipulativeness (Blair, 2007; Cleckley, 1941; Hare & Vertommen, 1991). Additionally, the Psychopathy Checklist-Revised Edition (PCL-R; Hare & Vertommen, 1991), generally viewed as a highly robust tool for measuring psychopathy, includes the previously mentioned traits plus superficial charm, pathological lying, failure to accept responsibility, need for stimulation, parasitic lifestyle, early behavior problems, lack of long term planning or goals, and promiscuous sexual behavior (Hare & Vertommen, 1991; Cooke & Michie, 2001). It is worth mentioning, at this point, that some debate remains about the central features of psychopathy as a construct. Measures, for example, that include traits such as boldness (e.g., Triarchic Psychopathy Measure (TriPM), Patrick, 2010) or the closely related fearless dominance (e.g., Psychopathic Personality Inventory – Revised (PPI-R), Lilienfeld & Widows, 2005) have received particular criticism (e.g., Miller & Lynam, 2012; but see Lilienfeld et al., 2012). Yet, it seems reasonable to suggest that the general consensus among scholars is that psychopathy represents a confluence of traits that predict a host of antisocial outcomes (Cooke & Michie, 2001; Hare, 1996; Patrick et al., 2006). Furthermore, measures, which include assessment of boldness/fearless dominance, have been widely used, including in research on the associations between psychopathy and intelligence, which will allow for an examination of (potential) differential associations between these traits.

Given the range of socially adverse outcomes often associated with psychopathy— including crime—it is perhaps tempting to conflate the construct with other well-established behavioral and personality disorders, antisocial personality disorder (ASPD) being chief among them. To be sure, there is a resemblance between the phenotypes. Yet, despite strong associations between them, and despite the fact that they predict similar outcomes, psychopathy and ASPD (and Conduct Disorder & Oppositional Defiant Disorder) are not fully interchangeable. As others have noted, in fact, when ASPD was added to the *Diagnostic and Statistical Manual for Mental Disorders – III* in 1980, the intent was that it would be a behavioral construct, which also captured variation in psychopathy, owing primarily to the belief that assessment of personality traits was fraught with measurement difficulties (see Hare, Hart, & Harpur, 1991). The result was a behaviorally based construct that poorly captured the nuances of psychopathy (Cooke & Logan, 2015; Hare, Hart, & Harpur, 1991). While scores on a measure of psychopathy have been found to correlate with symptoms of ASPD in prisoners (Hare, 2003), those labeled as psychopathic based off of diagnostic cut-offs on a psychopathy measure make up only a small subset of those who meet the diagnostic criteria for ASPD (Widiger, 2006).

At this point, it is useful to insert another key caveat; despite the prevalence and usefulness of research referring to “psychopaths” or individuals “with psychopathy,” taxonomic analyses have strongly suggested that psychopathy is best conceived of as a dimensional rather than categorical construct (e.g., Edens et al., 2006; Guay et al., 2007). This aligns with the growing support for a dimensional approach to personality psychopathology in general (e.g., Hopwood et al., 2018), and indeed DSM-5 now includes a dimensional Alternative Model for Personality Disorders (AMPD) in Section III. Although not included in the current iteration of the DSM, there is an ongoing effort within clinical psychology to create a model of psychopathology based on similar ideas to those undergirding the AMDP. Namely, that all of psychopathology is dimensional, rather than categorical, and that there is significant overlap and comorbidity between the categorical disorders currently in use (Kotov et al., 2017).

The key point is that regardless of whether they are treated categorically or dimensionally, psychopathic traits are clearly related to what the DSM labels ASPD (and Conduct Disorder (CD) and Oppositional Defiant Disorder (ODD)). Research using both categorical and dimensional approaches to assessing ASPD, for example, has found significant associations with psychopathic traits (e.g., Anderson et al., 2014; Few et al., 2015; Kavish, Sellbom, & Anderson, 2018). Furthermore, both psychopathy (e.g., Dolan & Anderson, 2002; Kavish, Bailey, Sharp, & Venta, 2018) and ASPD (e.g., Stevens, Kaplan, & Hesselbrock, 2003) have been associated with intelligence, as well as with overt behavioral problems. Yet, as previously noted, not all individuals who meet the criteria for disorders like ASPD necessarily score in the highest ranges on psychopathy measures (Widiger, 2006). Furthermore, psychopathy measures are typically less behaviorally saturated than the diagnostic criteria for ASPD and related disorders, and traits such as superficial charm or a lack of empathy are common to psychopathy measures, but absent from the assessment of ASPD. Therefore, we feel it is important to understand not only the associations between intelligence and psychopathic traits, but also the shared and unique associations between intelligence and constructs highly related to psychopathy, including ASPD, CD, and ODD— all of which we discuss in more detail in sections that follow below.

Put another way, although psychopathy, ASPD, CD, and ODD are not simply different names for the same construct, they can reasonably be considered as belonging to the same antisocial family of traits, and therefore, greater understanding is needed of what features (i.e., intelligence) might contribute to their overlap and distinctness. For ease of presentation, we will use the description “antisocial disorders” when referring to ASPD, CD, and ODD collectively, but we analyze them both collectively and separately so comparisons across literatures can be made.

## Intelligence

General intelligence, commonly referred to as *g* or the positive manifold, is arguably the best measured trait in all of psychology and research from a variety of disciplines has repeatedly found that it is immensely important in most areas of life (Gottfredson, 2002; Ritchie, 2015). Researchers have been studying and refining the concept of *g* since Spearman (1904) first proposed it in the beginning of the 20^th^ century as a way to conceptualize overall mental ability rather than variation across a specific type of ability (e.g. verbal ability or mathematical skill) (Gottfredson, 1997; 2002).

Similar to psychopathy, intelligence has consistently been linked to important life outcomes. IQ scores, which are meant to estimate general intelligence, predict socioeconomic status (Strenze, 2007), educational achievement (Deary, Strand, Smith, & Fernandes, 2007; Gottfredson, 1997; Lynn & Mikk, 2009; Strenze, 2007), occupational status and job success (Gottfredson, 2002; Strenze, 2007), mating success (Greengross & Miller, 2011), physical and mental health (Batty, Der, Macintyre, & Deary, 2006; Deary, Weiss, & Batty, 2010; Der, Batty, & Deary, 2009; Gottfredson & Deary, 2004) and longevity (Beaver et al., 2016; Deary, Weiss, & Batty, 2010; Gottfredson & Deary, 2004). Specifically, having a higher degree of intelligence has been found to be a predictor of completing more years of education (Deary, Strand, Smith, & Fernandes, 2007; Gottfredson, 1997; Lynn & Mikk, 2009; Strenze, 2007), gaining a higher status career (Gottfredson, 2002; Strenze, 2007) and living longer (Deary, Weiss, & Batty, 2010; Gottfredson & Deary, 2004). At the macro level, estimates of the mean IQ of a state (Kanazawa, 2006) or country (Jones, 2015) also predict differences in per capita Gross State Product (GSP) and Gross Domestic Product (GDP), respectively.

## Intelligence and Psychopathy

In *The Mask of Sanity*, Cleckley (1941) provided some of the earliest clinically based descriptions of psychopathy. One of the key attributes included in this description was that ‘psychopaths’ possess “good intelligence” (Cleckley, 1941). Since that early description, the conceptualization of psychopathy, especially in the public eye, has often depicted the “psychopath” as an evil genius or criminal mastermind (think Hannibal Lecter from *The Silence of the Lambs*; as described by DeLisi, Vaughn, Beaver, & Wright, 2010; Kavish, Bailey, Sharp, & Venta, 2018).

When considering the outcomes associated with intelligence and antisocial disorders, however, it seems reasonable to suggest that there might actually be a negative relationship between the two. For example, one of the largest behavioral overlaps between psychopathy and *low* intelligence is the increased propensity toward violent and criminal involvement. Numerous studies and reviews have found a robust negative relationship between intelligence and delinquency in adolescents and juveniles (Hernstein & C. Murray, 1994; Hirschi & Hindelang, 1977; Wilson & Hernstein, 1985). This relationship between intelligence and antisocial behavior continues into adulthood with lower intelligence scores being a significant risk factor for criminal behavior (Hernstein & C. Murray, 1994; J. Murray et al., 2010). Lower levels of intelligence have also been found to predict longer criminal careers (Piquero & White, 2003) and higher rates of violence among incarcerated individuals (Diamond, Morris, & Barnes, 2012). And on the opposite end of the spectrum, a meta-analysis of intelligence and crime found that higher intelligence was a protective factor against offending (Ttofi et al., 2016).

Similarly, psychopathy has been repeatedly associated with antisocial behavior and criminal activity (Hemphill, Hare, & Wong, 1998; Porter, Brinke, & Wilson, 2009; Salekin, Rogers, & Sewell, 1996). A meta-analysis of 53 studies totaling over 10,000 participants reported that psychopathy was a significant predictor of juvenile delinquency and assessment of psychopathy as a predictor of violence was found to be valid as early as middle childhood (Asscher et al., 2011). Additionally, psychopathic individuals tend to commit more violent crime (Porter, Brinke, & Wilson, 2009), more violence in prison (Hare, 1999), and recidivate at much higher rates (Hemphill, Hare, & Wong, 1998; Langevin & Curnoe, 2011).

Given the overlap in outcomes that correlate with both lower intelligence and psychopathy, researchers have more recently become interested in directly testing the link between the two phenotypes. The findings of this line of research have been relatively equivocal with many non-significant results (Dolan & Park, 2002; Hare & Jutai, 1988; Pham, Philippot, & Rime, 2000), as well as significant negative relationships (Dolan & Anderson, 2002). The ambiguity of these results and the common limitation of very small samples necessitate review and meta-analysis to further elucidate the possible link between intelligence and psychopathy at the construct level.

One meta-analysis has already examined the possibility of a relationship between psychopathy and intelligence and found no association (O’Boyle, Forsyth, Banks, & Story, 2013). However, the researchers were specifically interested in a Dark Triad (DT; Paulhaus & Williams, 2002) perspective, and consequently made inclusion/exclusion criteria decisions that left important gaps. Because the DT, the idea that so-called “dark traits” (psychopathy, narcissism, and Machiavellianism) are overlapping, yet distinct, traits that frequently co-occur, was conceptualized specifically for non-clinical populations, O’Boyle and colleagues (2013) excluded psychiatric, child, and incarcerated samples. Psychopathy is a personality construct with operationalizations (e.g., Cleckley, 1941) that predate considerably the DT, and it has been conceptualized and frequently assessed in all of these populations (e.g., Barry et al., 2000; Hare, 1991; 2003; Skeem & Mulvey, 2001). Indeed, one of the most popular measures of psychopathy, the Hare Psychopathy Checklist (PCL; Hare, 1991), was designed for use with prisoners. Thus, the decision to exclude these samples has left important questions unanswered, despite the results presented in O’Boyle et al. (2013).

The previous meta-analysis was also limited by its exclusion of a large number of studies which ultimately made examination of facet and factor level relationships impossible. Psychopathy is increasingly recognized as a multidimensional construct (e.g., Lilienfeld, 2018) and researchers have begun to attempt to further untangle the relationship between intelligence and psychopathy by evaluating how they are associated at the facet and trait level (e.g. DeLisi et al, 2010; Salekin, Neumann, Leistico, & Zalot, 2004). Preliminary results from this budding area of research suggest that particular facets of intelligence and factors of psychopathy could be driving the relationship that has been found in some of the construct level research. For example, verbal intelligence has been particularly implicated as being *positively* related to the interpersonal factor of psychopathy (Salekin et al, 2004) and negatively related to the affective (Salekin et al, 2004, Vitacco, Neumann, & Jackson, 2004) and behavioral/lifestyle components (Vitacco, Neumann, & Jackson, 2004). Moreover, there are numerous other variables that may moderate the relationship between facet and full-scale IQ and the factors and overall psychopathy (e.g., age, race, sex). Previous work (O’Boyle et al., 2013), examined only four potential moderators (age, sex, sample type, and measure of intelligence), leaving other potential moderators (e.g., measure of psychopathy, published vs unpublished data) untested. Finally, the potential problem of publication bias or low evidentiary quality in the literature was generally unaddressed.

## Overlap and Distinction Between Antisocial Disorders

As we’ve already discussed briefly at an earlier point, there exist multiple diagnoses for various presentations of problematic or antisocial behaviour. We first broached this issue by discussing ASPD, owing to its severity and close overlay with psychopathic tendencies. Also relevant in the current context, yet residing at the milder end of the spectrum, Oppositional Defiant Disorder refers to a pattern of argumentative and defiant behaviour (e.g., arguing with authority figures and refusing to comply with authority figures and rules), vindictiveness, and an angry or irritable temperament, typically emerging no later than adolescence. ODD is often a precursor to CD, which refers to a more concerning pattern of repetitive and persistent violation of societal norms and the rights of others, possibly including aggression towards people or animals (e.g., bullying or carrying a weapon), destruction of property, deceitfulness or theft, and serious rule violations (e.g., truancy). CD typically emerges in middle childhood or adolescence and is associated with an elevated risk for criminal behaviour and a diagnosis of ASPD as an adult. Antisocial Personality Disorder is an adult-disorder which requires evidence of CD prior to age 15, and reflects a pervasive pattern of disregard for and violation of the rights of others, often including criminal behaviour, impulsivity and recklessness, deceitfulness, and a lack of remorse (*DSM-5*).

Most importantly, all three disorders are associated with psychopathic traits (e.g., Kavish, Sellbom, & Anderson, 2018; Rogers e al., 1997; Salekin, Rogers, & Machin, 2001). Indeed, the current iteration of the DSM now includes specifiers for CD (i.e., Callous-unemotional traits) and for ASPD (i.e., psychopathy specifier) that are intended to more fully capture psychopathy. Quantitative genetics research has also found substantial genetic covariation between these constructs (e.g., between ODD and CD; Tuvblad et al., 2009), implying a similar, but not identical genetic etiology. As such, it may seem reasonable to expect these constructs to demonstrate somewhat similar associations with intelligence. Furthermore, since these constructs are typically related to poor decision-making, rule violations and/or criminal offending, and are typically seen as maladaptive, it might be expected that if these constructs, are inversely correlated with higher intellectual functioning (in contrast to the findings of O’Boyle et al, 2013).

Although these disorders typically covary, they are of course not perfectly correlated. Tuvblad and colleagues (2009) found the genetic covariation between CD and ODD to be approximately 0.43, the shared environmental correlation to be approximately 0.77, and the non-shared environmental correlation to be about 0.22. Moreover, while it seems reasonable to generally hypothesize a negative relationship between intelligence and each of these disorders resulting from the consistent associations found between low IQ and antisocial behaviour (e.g., Diamond, Morris, & Barnes, 2012; Piquero & White, 2003), it is plausible that the *behavioral constructs*, particularly CD and ASPD, will evince the strongest negative associations with intelligence. Psychopathy, as a measure rooted more in personality-based traits, would then be expected to demonstrate the weakest association as the associations between intelligence and personality traits (e.g., the Five Factor Model) are typically modest (Bartels et al., 2012).

## Study Aims

With the above in mind, there remain several gaps in the literature on the relationship between psychopathy, antisocial disorders and intelligence that we seek to fill. First, we seek to provide an estimate of the overall magnitude of the association between intelligence and psychopathy that is derived using less restrictive inclusion criteria, and that more deeply explores potential moderators and publication bias. Due to the multidimensionality of psychopathy (Lilienfeld, 2018), we will also explore the association between intelligence and psychopathy in finer detail by analyzing associations at the factor (Factor 1 and Factor 2) and facet (affective, interpersonal, lifestyle, and antisocial; or Boldness/Fearless Dominance and Impulsive-antisociality/Meanness and Disinhibition) levels of psychopathy. Similarly, measures of intelligence typically assess multiple domains of cognitive ability, which, although they are highly correlated and load onto the general factor of intelligence, are not perfectly associated and may be differentially associated with other variables. Therefore, we also examine the associations of verbal intelligence (VIQ) and performance intelligence (PIQ) with antisocial disorders (and their facets) wherever possible.

Second, we seek to examine the association between intelligence and antisocial disorders closely tied to psychopathy. Because all three constructs (ASPD, CD, and ODD) are closely related to each other and, to an extent, meant to capture the same underlying theme (i.e., problematic behaviour and associated personality traits), we first consider the association between intelligence and the antisocial constructs combined. Yet, as we previously noted, these constructs diverge in the specific behaviours and traits by which they are operationally defined. Thus, we also consider the link between each disorder and intelligence individually to assess the possibility that there are differential associations (e.g., perhaps ASPD as a more behavioral construct will be particularly associated with intelligence compared to the broader behaviorally and personality influenced construct of psychopathy).

### Moderators

In addition to considering overall (and domain/factor/facet level) associations between intelligence and our four antisocial disorders (psychopathy, ASPD, CD, & ODD), we seek to understand how the associations we find might be moderated by other factors. Specifically, we will assess whether the associations are moderated by sample type, geographic location, age group, sex composition, instruments used to assess antisocial disorders, instrument used to assess IQ, whether or not a given study included covariates, publication status, and publication year. The sample type (clinical, forensic, or community) will be included as a moderator because prior research on psychopathic traits using the Self-Report Psychopathy Scale – Short Form (Paulhus et al., 2016) failed to find measurement invariance between incarcerated and student samples (Debowska et al., 2018), suggesting psychopathic traits may manifest differently across samples.

The specific measures used to assess psychopathic traits (or antisocial disorder) and intelligence will also be assessed as potential moderators because, although psychopathy measures (for example) typically correlate, they do not perfectly correlate and differ in the extent to which they capture various features (e.g., antisocial behaviour and boldness). If the measure used to assess psychopathy (or ASPD, CD, or ODD) is found to be a significant moderator, this would suggest that the way in which a particular construct is measured informs whether, and to what extent, it is associated with intelligence. In other words, if the PCL family of psychopathy measures is found to be associated with intelligence measures (while other measures of psychopathy are not), this could suggest that antisocial behaviour (more explicitly assessed by the PCL) is primarily driving psychopathy’s association with intelligence. Alternatively, it could suggest that traits such as boldness (more explicitly assessed by measures such as the TriPM and PPI-R) are related to intelligence in the opposite direction of other psychopathic traits, and thus, work to reduce the magnitude of the association between global psychopathy scores and intelligence. Publication status (published or unpublished), moreover, could moderate the effect size because research shows that significant results are more likely to be submitted and accepted for publication, contributing to the well-known “file drawer effect” in which non-significant results do not enter the knowledge base for a particular topic (Rosenthal, 1979).

A final moderator of interest is the gender of the participants in the samples (all males, all females, or mixed). To date, it remains unclear whether gender may moderate the association between antisocial disorders and intelligence, though there are some reasons to expect as much. First, there are clear and pronounced gender differences in antisocial disorders. For example, Cale and Lilienfeld (2002) reviewed the existing literature and concluded that the prevalence of categorically assessed ASPD is higher in males and that males score higher than females on dimensional measures of psychopathy and ASPD. The review also stated that there is tentative evidence for gender differences in the neuropsychological correlates of ASPD. Cale and Lilienfeld (2002) further reported evidence that the developmental trajectory of psychopathy and ASPD differs between males and females, including that males demonstrate more externalizing and less internalizing behaviours between age five and adolescence, and that gender differences in the types of aggression and antisocial behaviour are exhibited across development and into adulthood.

Looking more broadly at sex differences in neuroanatomical and neurological functioning, Ritchie and colleagues (2018) reported higher raw volumes, raw surface areas, and white matter fractional anisotropy in males and higher raw cortical thickness and white matter tract complexity in females. The authors further reported connectome differences including stronger connectivity in unimodal sensorimotor cortices in males and stronger connectivity in the default mode network for females. As the authors also note, these subregional differences were not accounted for by differences in broader variables such as total volume, total surface area, average cortical thickness, or height. Of particular importance for the consideration of sex as a moderator in the current study, however, sex differences in brain volume and surface area were associated with sex differences in verbal-numerical reasoning, and differences in structure and connectivity were also found in subregions of the brain associated with emotion and decision making. In sum, then, there appears to be some evidence for sex differences in psychopathic and other antisocial traits (e.g., Cale & Lilienfeld, 2002), some evidence for modest differences in cognitive abilities (e.g., Ritchie et al., 2018), and evidence for differences in neuroanatomical features that are correlated to emotions and decision making (e.g., Ritchie et al., 2018), which are relevant variables to psychopathy and antisocial behaviour.

## Method

### Inclusion criteria

Although this meta-analysis was not preregistered, studies included in the meta-analysis had to meet the following inclusion criteria:

1. The study measured the association between intelligence and a measure of psychopathy and/or antisocial personality traits.
2. The study examines at least one component of psychopathy (i.e. Factor 1, Factor 2, and/or one of its facets). For studies reporting the Dark Triad, only the psychopathy scale was included (for other dimensions see ÓBoyle, Forsyth, Banks & Story, 2013). With regard to the antisocial personality criteria, the study had to define antisocial behaviour in terms of psychiatric diagnoses based on psychiatric manual such as *Diagnostic and Statistical Manual of Mental Disorders (any version)* (DSM, The American Psychiatric Association) or *International Statistical Classification of Diseases 10^th^ Revision* (ICD-10, WHO, 1992) [i.e., oppositional defiant disorder (ODD), conduct disorder (CD), disruptive behaviour disorder (DBD), antisocial personality disorder (ASPD)]. If the antisocial behaviour was exclusively based on legal operationalization (i.e. reported delinquency, number of convictions and/or criminality, or aggression/violence) the study was excluded. Studies that used antisocial behaviour questionnaires associated with the DSM (i.e., CBCL, Strengths and Difficulties Questionnaire (SDQ; Goodman, 1997) or psychopathy measures (e.g., Antisocial Behaviour Questionnaire (ABQ; Loeber, Stouthamer-Loeber, Van Kammern, & Farrington, 1989) were included (see Fontaine, Barker, Salekin & Viding, 2008; Rispens et al. 1997; Wall, Sellbom & Goodwin, 2013). Other questionnaires must be supplemented with psychiatric diagnostic criteria. Four studies (Ford, Farah, Shera, & Hurt, 2007; Hofvander et al. 2011; Masten et al. 1999; and Nomura, Rajendran, Brooks-Gunn & Newcorn, 2008) used questionnaires without psychiatric diagnosis (e.g., the Life History of Aggression scale; Cocarro et al., 1997). They were included because the items were similar to those studies with psychiatric diagnostic criteria (see sensitivity analysis). Samples with ADHD were excluded given the existence of different background factors and correlates (Lynam, 1996). Finally, studies using the five-factor model (FFM) to measure antisocial personality traits were excluded (for review, see Decuyper, de Pauw, de Fruyt, de Bolle & de Clercq, 2009).
3. The study had to include a standardized intelligence test. The Wechsler tests remain widely utilized, but any other standardized test of IQ was included. We included any version, abbreviated versions, and any subscale of the WAIS (i.e. Vocabulary, Digit Symbol). Verbal IQ (VIQ; broadly the ability to analyse information and solve problems using language and language-based reasoning skills) and Performance IQ (PIQ; reflects non-verbal, visuo-spatial abilities) were recorded when available. We excluded the PIQ–VIQ discrepancy (for review, see Isen, 2010). Although, measures of working memory do correlate with intelligence these cognitive tests were excluded. Additionally, we excluded emotional intelligence measures.
4. The study had to report the zero-order correlation or the necessary data to calculate the zero-order correlation between IQ and psychopathic or antisocial personality traits. When effect sizes were reported that controlled for covariates (i.e. studies that reported effect sizes as coefficients in a regression with multiple variables entered simultaneously in the model, partial correlations, or structural equation models), the authors were contacted to request the data for zero-order correlations. The studies with covariates were coded to be included in moderator analyses.
5. No restrictions were applied on the following categories: type of population (clinical, institutional and general population), age (adults, adolescents and children) and gender (males, females and mixed).

### Literature search strategies

To identify studies for inclusion in the meta-analysis, we conducted searches in the following bibliographic databases: MEDLINE, Web of Knowledge, SCOPUS and Google Scholar. We limited the search to peer-reviewed studies published in English between 1940 and the most recent search date (May 2017) because the first major operationalization of psychopathy appeared in 1941 (Cleckley, 1941). The search for candidate studies to be included in the meta-analysis was conducted using keywords relevant to (“*antisocial personality disorder*” OR “*psychopathy*” OR “*conduct disorder”* OR *“oppositional defiant disorder (ODD)”* OR *“disruptive behavior disorder (DBD)”)* AND (“*IQ*” OR “*intelligence”* OR “*general cognitive abilities”)* AND *(“community”* OR *“students”* OR *“offenders”* OR “*adolescents*”). The search terms were used separately and in different combinations for the database searches. In total 17 personality terms were included regarding antisocial traits (e.g., psychopath*, psychopathic traits, callous-unemotional trait, antisocial personality) and measures (e.g., PCL, LSRP, PPI, DSM). Sixteen general cognitive abilities terms were used (e.g., intellectual functioning, intellectual abilities, general cognitive functioning) and their measures (e.g., WAIS; NART; SILS, Raveńs Progressive Matrices) and 12 population type terms (e.g., youth, juvenile, criminal, offender, viol*, student, general population, community, students, children). Additional articles were obtained through inspection of the reference lists of articles and reviews obtained in the above search. Finally, we contacted authors for unpublished data on the topic under investigation.

### Coding moderators

Study characteristics were coded by the first author using a coding form (when any questions arose, NK and BBB consulted). The following moderators were coded for each correlation, with the coding rules based on previous meta-analyses (see Decuyper et al. 2009; Isen, 2010).

#### Age group of participants

Age groups were coded as follows: child (1–11 years old), adolescent (12–17 years old), or adult (18 years old or older) (Lorber, 2004). If studies considered age ranges, the mean age of the sample was included. Furthermore, age was not reported in some studies of incarcerated individuals or college undergraduates. In these cases, a code of “adult” was assigned (college students tend to be within 18–22 years of age) (Isen, 2010).

#### Gender

A sample was coded as “males”, “females” or “mixed”. In mixed samples, the percentage of males in the sample was calculated.

#### Outcome type

Antisocial behavior type was coded as one of three categories based on a clear definition given by the DSM or specific questionnaire or interview based on the diagnostic manual (the children and adolescent disorders (CD, ODD) and the adult disorder, ASPD). Psychopathy was coded as derived from Hare’s/Cleckley’s psychopathy description or other validated models (i.e. triarchic (Tri) construct and Dark Triad). In studies in which ASPD and psychopathy were both reported, effect sizes were classified based on the measure reported in the paper.

#### Psychopathy/ASPD measure

All versions of Harés Checklist for psychopathy were considered as the category “PCL”. Other inventories or questionnaires of psychopathy, conduct disorder, and antisocial personality traits were coded as “Other inventory”, and the category “Interview” was used for the DSM or any other type interview used to measure antisocial personality traits.

#### IQ measure

All versions of the WAIS and WISC were collapsed into a single category “Wechsler” (i.e., WISC, WAIS, WASI, WIP, etc.). The rest of the tests were coded as “Other” (Isen, 2010).

#### Recruitment source

Recruitment was coded as “Clinical” (i.e., university evaluation units; referral clinics and courts; social services agencies; psychiatric hospitals and assessment units), “Institutional” (i.e., prisons, youth detention centers, security hospitals and probationary supervision) or “Community” (i.e. schools, university, general population, Navy Center, prenatal clinics and birth cohort).

#### Covariates

As pointed above, some studies reported regression models or adjusted correlations. This was coded as “Yes/No”.

#### Region

Region of each sample was coded as “North American”, “European” and “Australia/New Zealand” (Decuyper et al. 2009).

#### Publication type

This was coded as “published data” and “unpublished data”. The latter was grey literature, that is, PhD Dissertations and data reported by authors via email, not reported in a published article.

### Selection and calculation of effect sizes

The collected data (mainly Pearson correlations) were analyzed using Fisher’s Z-transformed correlation coefficients weighted by the inverse of the Variance (see Lipsey & Wilson, 2001) using Comprehensive Meta-Analysis (CMA) version 3.3 statistical software (Borenstein, Hedges, Higgins, & Rothstein, 2007) in combination with ‘lavaan’, a meta-analysis package for R statistical computing environment (R Core Team, 2013). A random effects model was applied to examine the overall association between intelligence and psychopathy and related antisocial disorders (see Lipsey & Wilson, 2001) because we assumed that effect sizes would vary across studies. Cohen (1988) suggested that the effect size (*r*s) of .10, .30 and .50 be considered small, medium and large, respectively. Yet it is worth noting that for psychological research, others have recommended interpreting the effect sizes of .10, .20 and .30 as small, moderate and large effects, respectively (Hemphill, 2003).

Homogeneity (*Q* and I^2^) tests were performed to determine whether the studies can reasonably be described as sharing a common effect size. That is, is the variation among study outcomes due to random chance (*Q t*est) and what percentage of variation across the studies is due to significant heterogeneity (I^2^ test) (see Hedges & Olkin, 1985, Higgins, Thompson, Deeks & Altman, 2003). Generally, I^2^ values of 25%, 50%, 75% represent low, moderate and high between-study heterogeneity (Q*_B_*) (Higgins et al., 2003), with higher levels of heterogeneity suggesting that greater proportions of between-study variation in effect size are due to differences between the studies (e.g., such as using different measures of intelligence). A low I^2^ value on the other hand, would indicate that between-study variation is due mostly to chance.

Thus, we can use these analyses to examine the role of potential moderator variables and determine if we need to conduct additional analyses. For all categorical variables, moderator analyses were conducted using the analog to the ANOVA (with random effects), whereas fixed effect meta-regression analyses were conducted for the continuous moderator variables (e.g., publication year). We also evaluated the within-group heterogeneity (Q*_W_*) for moderator variables. Significant values within these analyses indicate that there is significant heterogeneity between studies *within* a moderator category.

Finally, we estimated the robustness of the meta-analytical estimates by performing sensitivity analyses. These adjust for the impact of publication bias as well as the impact of outliers and influential studies. We applied the trim-and-fill method (Duval & Tweedie, 2000) to identify and adjust for publication bias and the Egger’s linear regression procedure (Sterne & Egger, 2001). When the relationship between IQ and an outcome variable was reported for multiple measures of the outcome variable, we selected the best known, most validated and reliable instrument that most closely operationalized the antisocial personality or psychopathy outcome of interest. When different IQ measures were reported for an outcome, we applied the same criteria. For studies reporting more than one effect size (e.g., more than one sample in a single study), we calculated the mean of the effect sizes along with the variance for the mean effect size. This weighted mean effect size was then included in our analyses.

When a study reported only Factors 1 and 2 or facet correlates, we averaged them to create a mean effect size. Composite scores were only created when all dimensions of the measure were available (ÓBoyle et al., 2013), that is, if only two facets were reported, we did not create a total psychopathy score. However, in follow up moderator analyses, we also calculated effect sizes for Factors 1 (interpersonal/affective or callous-unemotional in youth) and 2 (lifestyle/antisocial) of psychopathy or facets when available in order to determine whether IQ was more strongly associated with a particular factor or facet. In addition, when VIQ and PIQ were reported, we averaged them, but also recorded them for moderator analyses. If the study reported the Vocabulary and/or Similarities subscales, it was included in the VIQ moderator analysis; and if the Block Design and/or Matrix Reasoning subscales were reported, they were included in the PIQ.

In order to statistically disentangle shared effects of the facets of intelligence on psychopathy, ASPD, & CD, we used meta-analytical structural equation modeling (i.e., MASEM) to produce relative weights (Johnson, 2000). Relative weights provide two coefficients. First, relative weights display the amount of variance explained in an outcome that is accounted for by the predictor (e.g., VIQ or PIQ). Second, rescaled relative weights reflect the percentage of variance accounted for by each predictor. Rescaled relative weights are determined by dividing the individual relative weights by the total model *R^2^* (LeBreton, Hargis, Griepentrog, Oswald, & Ployhart, 2007).

In order to conduct MASEM, a three-step process was conducted for each outcome (i.e., psychopathy, ASPD, & CD) per the practices outlined by Viswesvaran & Ones (1995):

1. zero-order meta-analytic relationships between VIQ, PIQ, and each outcome included in this study were calculated as described above. Because the inter-correlation between VIQ and PIQ was not produced in this study but is necessary for determining the relative contribution of each predictor, the zero-order meta-analytic inter-correlation was taken from Isen (2010).
2. Individual correlation matrices were constructed for each outcome such that a 3×3 matrix included correlations between VIQ, PIQ, and an outcome. (3) Each correlation matrix was used in path-analysis using ordinary least squares regression (Viswesvaran & Ones, 1995). Furthermore, the sample size for each path model was determined by the harmonic mean of the sample sizes across the correlations within each path meta-analytic correlation table.

The PCL was the most commonly used psychopathy measure across the studies, so we collapsed the different components of other measures (i.e., subscales of MPQ-Tri, PPI, PAI-ANT) into PCL-Factor 1 (Interpersonal and Affective) and 2 (Antisocial Behavior and Impulsivity). Our criteria were based on previous correlational studies (see Benning, Patrick, Blonigen, Hicks, & Iacono, 2005; Brislin, Drislane, Smith, Edens & Patrick, 2015, Copestake, Gray & Snowden, 2011; Venales, Hall & Patrick, 2014). In particular, Factor 1 of the PCL-R consisted of the concept of meanness and boldness on the TriPM, and Fearless Dominance and Coldheartedness on the PPI-R, whilst Factor 2 was represented by Disinhibition on the TriPM and Impulsive-Antisociality on the PPI-R. Some studies reported overall correlations, as well as separate correlations for males and females; or children, adolescents and adults. When this occurred, independent effect sizes for each one were included in order to use these effect sizes for the gender and age moderator analyses.

### Transparency

As we have mentioned, this meta-analysis was not preregistered. Sample sizes were derived directly from the published material or reported by original authors, if unpublished. The CMA software used for this meta-analysis is operated via a “point-and-click” interface and thus does not provide a method for recording/extracting syntax, so we provide full tables of information entered for each study in the paper which can be accessed on the Open Science Framework (https://osf.io/7uvcp/). While we are unable to provide a complete analysis script, script is available (on the OSF project page for this paper at the link above) for the MASEM portion of the anlayses, as those models were estimated using the software package R. Because the current study is a meta-analysis, most of the included data is already publicly available (i.e., published manuscripts), but again all data is reported on the Open Science Framework in files available at the above link. We report test statistics, p-values (in-text or in tables), and confidence intervals for all analyses.

## Results

A detailed list of the studies included in the meta-analysis is provided in Supplementary Table 1S (https://osf.io/7uvcp/; see Figure 1 for further information regarding excluded studies). The final sample consisted of 94 studies, reporting 143 total correlations, published during the period of 1965 to 2017. Data were obtained from a total of 46,784 subjects, comprising independent samples. Nearly all studies were conducted in the United States, but other nationalities were represented such as England, Germany, Canada, New Zealand, Australia, Netherlands, Finland, Sweden, France, Romania, Italy, Malaysia, Bulgaria, Switzerland, and Spain.

**Figure 1.**
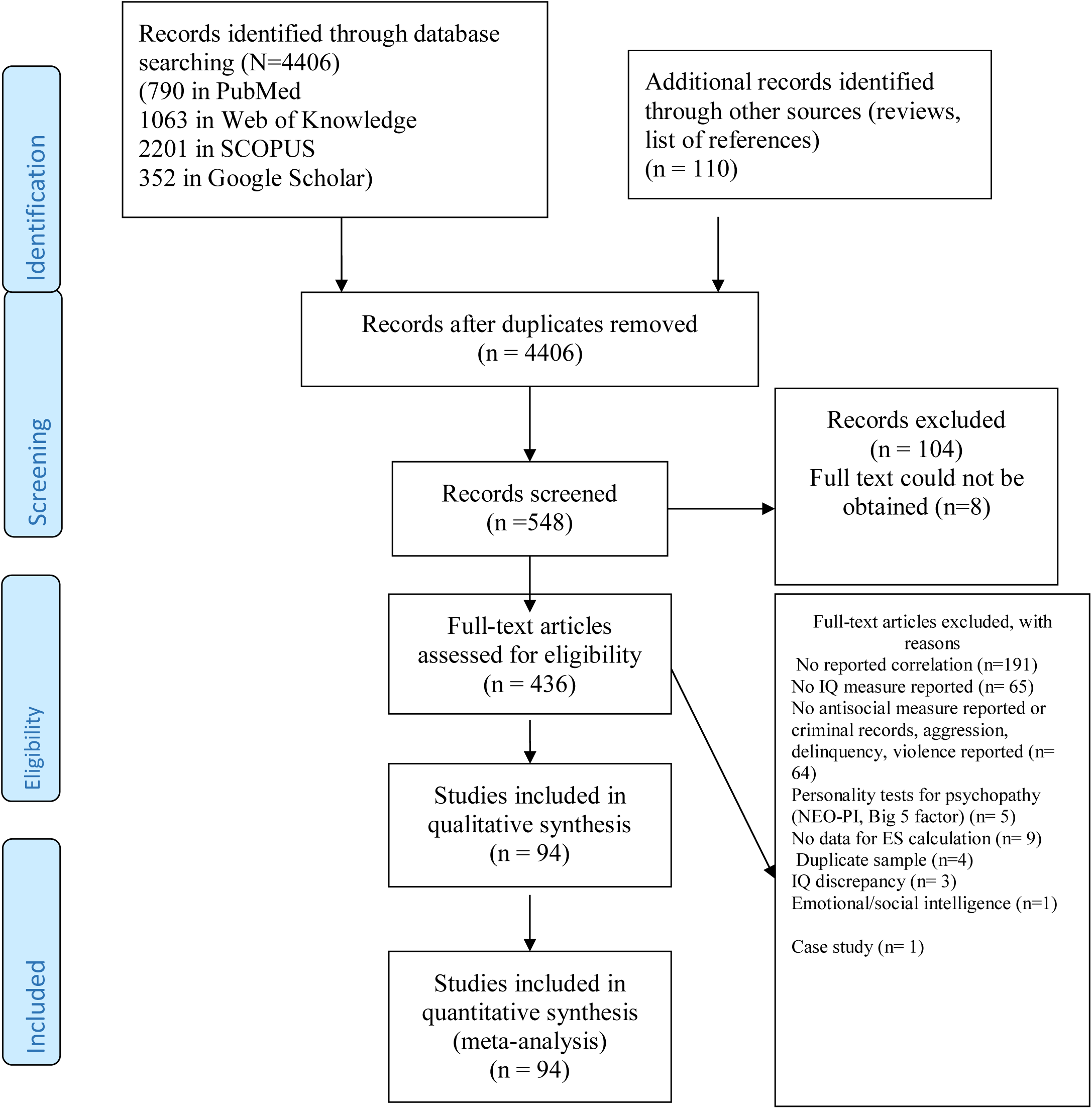
PRISMA Flow Diagram^1^.

### Intelligence and Psychopathy

Meta-analytic effect sizes for the associations between intelligence (FSIQ, VIQ, and PIQ) and psychopathy (psychopathy, Factor 1 psychopathy, and Factor 2 psychopathy) are presented in Table 1. Across 105 correlations, a weak but significant negative association emerged between FSIQ and psychopathy (r= -.07, p =.000). At a finer grained level of intelligence, VIQ was weakly associated (*r*= -.12, p= .000) with psychopathy, and a weak relationship was found for psychopathy and PIQ (r= -.05, p= .016). When examining factors of psychopathy, the meta-analysis for FSIQ and Factors 1 and 2 of psychopathy revealed a non-significant relationship (r= .01, p= .797) and a weak, significant, negative relationship (r= -.09, p= .000), respectively. Thus, there appears to be no association between FSIQ and Factor 1, which encompasses the interpersonal and affective component of psychopathy. There does, however, appear to be a small negative association between FSIQ and Factor 2 psychopathy, suggesting that those who score higher on the impulsive and antisocial behavior component of psychopathy tend to score lower on intelligence tests. Put another way, the association between intelligence and psychopathy seems to be driven by psychopathy measures, or domains of such measures, that index risky and antisocial behavior. This finding aligns with prior work on the association between indicators of intelligence and overt forms of aggression (e.g., Duran-Bonavila et al., 2017; Kennedy et al., 2011).

**Table 1.**
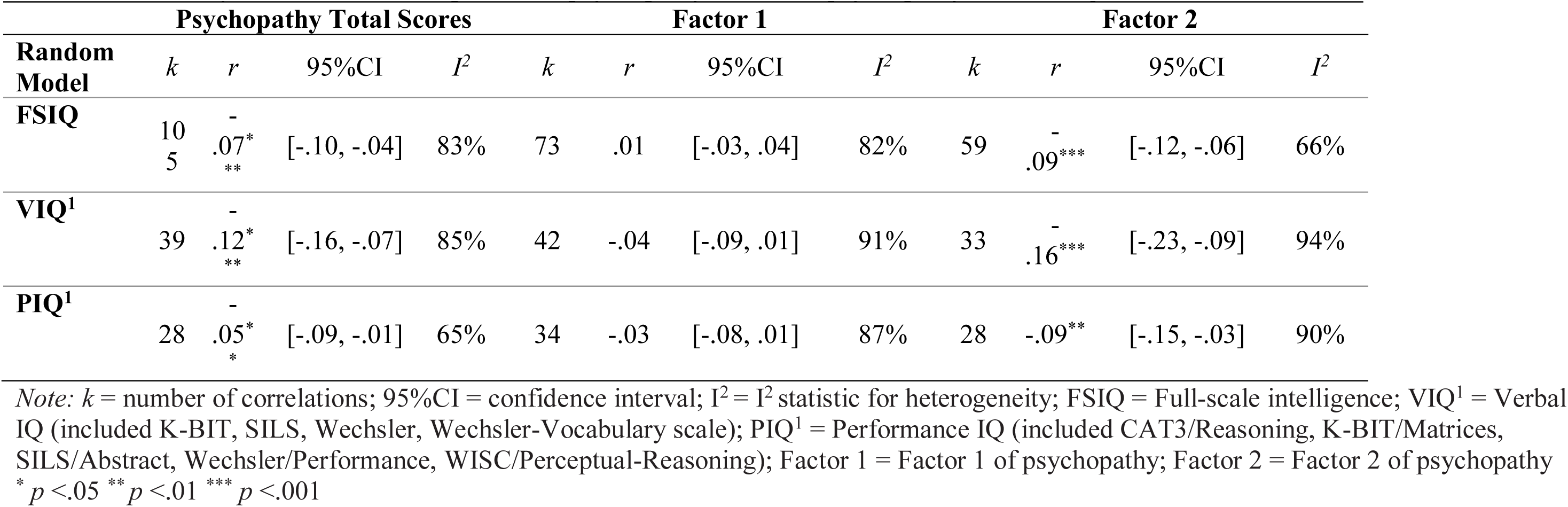
Meta-analysis of the relationship between psychopathy, factors of psychopathy, and intelligence

In keeping with our efforts to include as fine-grained an evaluation of these associations as possible, we next tested the associations between Factors 1 and 2 of psychopathy and both VIQ and PIQ (Table 1). When examining facets of intelligence, Factor 1 demonstrated a non-significant relationship with VIQ (r= -.04, p= .118) and PIQ (r= -.03, p= .169). With regard to Factor 2 psychopathy, analyses yielded a small negative relationship with VIQ (r= -.16, p= .000), and a relatively small negative relationship with PIQ (r= -.08, p=.011). In other words, no association was found between Factor 1 psychopathy and either domain of intelligence, but lower VIQ and PIQ were each associated with higher scores on the second factor of psychopathy.

Additionally, we analyzed the relationship between *facets* of psychopathy (1: Interpersonal; 2: Affective; 3: Lifestyle; and 4: Antisocial) and intelligence (FSIQ, VIQ, PIQ) (see Table 2). The meta-analyses revealed that the interpersonal facet 1 showed a significant small effect size for FSIQ that was positive (*r* = .14, p= 0.044). The correlations between facet 1 and VIQ and PIQ were negative for VIQ (*r* = -.03, p= 0.456) and positive for PIQ (*r* =.05, p= 0.756), yet neither was statistically significant. The affective facet 2 demonstrated a significant small negative effect size for all types of intelligence (FSIQ: *r* = -.16, p= 0.003; VIQ: *r* = -.18, p= 0.000; PIQ: *r* = -.12, p= 0.000). The lifestyle facet 3 revealed significant small negative effect sizes for VIQ (*r* = -.19, p= 0.000) and PIQ (*r* = -.17, p= 0.000), but a non-significant negative correlation for FSIQ (*r* = -.05, p= 0.371). Finally, the antisocial facet 4 showed a negative effect size for both VIQ (*r* = -.18, p< 0.001) and PIQ (*r* = -.20, p< 0.001), but a non-significant negative effect size for FSIQ (*r* = -.12, p= 0.278).

**Table 2:** Meta-analysis of the relationship between facets of psychopathy and intelligence

| Random Model | Facet 1 (Interpersonal) |  |  |  | Facet 2 (Affective) |  |  |  | Facet 3 (Lifestyle) |  |  |  | Facet 4 (Antisocial) |  |  |  |
| --- | --- | --- | --- | --- | --- | --- | --- | --- | --- | --- | --- | --- | --- | --- | --- | --- |
|  | k | r | 95%CI | I<sup>2</sup> | k | r | 95%CI | I<sup>2</sup> | k | r | 95%CI | I<sup>2</sup> | k | r | 95%CI | I<sup>2</sup> |
| FSIQ | 11 | .14* | [.00, .28] | 87% | 12 | -.16** | [-.27, -.05] | 83% | 11 | -.05 | [-.17, .06] | 81% | 6 | -.12 | [-.33, .10] | 85% |
| VIQ | 11 | -.03 | [-.11, .05] | 89% | 10 | - .18*** | [-.26, -.10] | 88% | 11 | - .19*<br>** | [-.28, -.11] | 92% | 9 | - .18*** | [-.25, -.10] | 90% |
| PIQ | 9 | .05 | [-.26, .35] | 99% | 8 | - .12*** | [-.18, -.07] | 53% | 9 | - .17*<br>** | [-.21, -.12] | 62% | 7 | -.20 | [-.47, .10] | 99% |

Note: *k* = number of correlations; 95%CI = confidence interval; I<sup>2</sup> = I<sup>2</sup> statistic for heterogeneity; FSIQ = Full-scale intelligence; VIQ = Verbal IQ; PIQ = Performance IQ. Facets included different measures (e.g., YPI), not only the PCL.
\* *p* < .05 \*\* *p* < .01 \*\*\* *p* < .001

As mentioned in our introduction, although controversial, some operationalizations of psychopathy have included a component related to emotional stability and social potency (i.e., boldness or fearless dominance). We conducted a final set of analyses for psychopathy (see Table 3) on measures such as the PPI and triarchic psychopathy scales (such as those created from items in the Multidimensional Personality Questionnaire, Tellegen, 1982; see Brislin et al., 2015; MPQ-Tri). When the Fearless Dominance (PPI) and Boldness (MPQ-Tri) subscales were considered together, the effect size for FSIQ was significant, small, and positive (*r* = .15, p= 0.002), but the effect sizes for the VIQ (*r* = .06, p= 0.311) and PIQ (*r* =.05, p= 0.244) were non-significant. Conversely, impulsive-antisociality (PPI) and triarchic meanness and disinhibition (MPQ-Tri) subscales were combined and yielded negative weak significant correlations with all types of intelligence (*r* = -.06, p= 0.050). We chose to combine PPI impulsive-antisociality with both TriPM Meanness and Disinhibition due to research suggesting that PPI impulsive-antisociality is strongly related to both domains of the TriPM (e.g., Drislane et al., 2014), as well as recent research on the factor structure of the original TriPM and the triarchic scales developed in other measures (e.g., MPQ-Tri) which found the Meanness and Disinhibition domains to be indistinguishable (Collison et al., 2019).

**Table 3.** : Meta-analysis of the relationship between PPI and MPQ-Tri subscales and intelligence

|  | **PPI-Fearless Dominance**<br>**and**<br>**MPQ-Tri-Boldness** |  |  |  | **PPI/Impulsive-Antisociality**<br>**and**<br>**MPQ-Tri/Disinhibition,**<br>**Meanness** |  |  |  |
| --- | --- | --- | --- | --- | --- | --- | --- | --- |
| **Random Model** | *k* | *r* | 95%CI | *I*<sup>2</sup> | *k* | *r* | 95%CI | *I*<sup>2</sup> |
| FSIQ | 6 | .15** | [.05, .24] | 72% | 6 | -.06* | [-.12, -.00] | 41% |
| VIQ | 4 | .06 | [-.06, .18] | 81% | 4 | -.06* | [-.10, -.00] | 7% |
| PIQ | 4 | .05 | [-.03, .13] | 60% | 4 | -.06* | [-.10, -.02] | 0% |

*Note:* *k* = number of correlations; 95%CI = confidence interval; *I*<sup>2</sup> = *I*<sup>2</sup> statistic for heterogeneity; FSIQ = Full-scale intelligence; VIQ = Verbal IQ; PIQ = Performance IQ.
\* *p* < .05 \*\* *p* < .01 \*\*\* *p* < .001

### Intelligence and other Antisocial Constructs

After meta-analyzing the various associations between intelligence and psychopathy, we next analyzed the relationships between intelligence (FSIQ, VIQ, and PIQ) with ASPD, CD, and ODD (Table 4). A total of 14 correlations between ASPD and FSIQ were included in the analysis, and the results revealed an overall significant, albeit small, negative correlation (*r=* -.13, p= .001). No significant association was found between ASPD and either VIQ (*r* = -.01, p= .770) or PIQ (*r* = -.01, p= .871), although these effect sizes were calculated from only four and three studies, respectively. The correlation for CD (*r=* -.11, p= .001) was based on analysis of 23 studies and was comparable to the association found for ASPD, indicating that lower intelligence was associated with higher scores on measures of both ASPD and CD. In contrast to ASPD, however, CD was negatively associated with both VIQ (*r* = -.17, p= .000) and PIQ (*r* = -.16, p= .025). Finally, with regards to FSIQ and ODD (*r=.06*, p= .001), a weak, positive association was uncovered. However, calculations for the relationship between intelligence and ODD should be interpreted with caution because they were based on only three correlations. Readers interested in results for ODD can access them in the supplementary material (Table 2S and Figure 1S), but given the low number of studies that were eligible for inclusion in our paper, ODD was excluded from the rest of the analyses we present here.

**Table 4.** Meta-analysis of the relationship between ASPD, CD, ODD and intelligence

| Random<br>Model | ASPD |  |  |  | CD |  |  |  | ODD |  |  |  |
| --- | --- | --- | --- | --- | --- | --- | --- | --- | --- | --- | --- | --- |
|  | *k* | *r* | 95%CI | *I*<sup>2</sup> | *k* | *r* | 95%CI | *I*<sup>2</sup> | *k* | *r* | 95%CI | *I*<sup>2</sup> |
| FSIQ | 14 | -.13*** | [-.20, -.06] | 74% | 23 | -.11*** | [-.18, -.04] | 95% | 3 | .06*** | [.04, .08] | 0% |
| VIQ<sup>1</sup> | 4 | -.01 | [-.09, .06] | 0% | 7 | -.17*** | [-.26, -.08] | 86% | 1 | .09 | [-.11, .28] | 0% |
| PIQ<sup>1</sup> | 3 | -.01 | [-.10, .08] | 13% | 4 | -.16* | [-.29, -.02] | 94% | - | - | - | - |

*Note.* FSIQ = Total score of intelligence; VIQ= Verbal score of intelligence; PIQ= Performance score of intelligence; k= number of correlations; 95%CI= confidence interval; *I*<sup>2</sup> = *I*<sup>2</sup> statistic for heterogeneity; ASPD = Antisocial Personality Disorder; CD = Conduct Disorder; ODD = Oppositional Defiant Disorder
\* p <.05 \*\* p <.01 \*\*\* p <.001

### Meta-analytic Structural Equation Modeling

Finally, we also examined the unique associations of VIQ and PIQ with our antisocial variables using meta-analytic structural equation modeling (see Table 5). As a set, VIQ and PIQ account for a statistically significant, yet small amount of variance in psychopathy (*R^2^* =.01) and CD (*R^2^* = .04), but not ASPD (*R*^2^ = .00). Disentangling the unique effects of each intelligence type, VIQ (91.255%) accounts for more variance in psychopathy than PIQ (8.745%). In CD, the two facets of intelligence account for more even proportions of the variance; with VIQ (54.173%) accounting for slightly more variance in CD than PIQ (45.827%).

**Table 5.** Results of meta-analytic SEM examining unique contributions of VIQ and PIQ

| Psychopathy | IQ Type | *B* | *SE<sub>B</sub>* | t-value | *p* | RW | %*R*<sup>2</sup> |
| --- | --- | --- | --- | --- | --- | --- | --- |
| *R*<sup>2</sup> = .01 | VIQ | -0.12 | 0.01 | -13.56 | <.001 | 0.013 | 91.255 |
| *F* = 111.20, *p* < .05 | PIQ | -0.01 | 0.01 | -0.59 | 0.55 | 0.001 | 8.745 |
| ASPD | IQ Type | *B* | *SE<sub>B</sub>* | t-value | *p* | RW | %*R*<sup>2</sup> |
| *R*<sup>2</sup> = .00 | VIQ | -0.01 | 0.03 | -0.21 | 0.83 | 0.000 | 50.000 |
| *F* = 0.07, *p* > .05 | PIQ | -0.01 | 0.03 | -0.21 | 0.83 | 0.000 | 50.000 |
| CD | IQ Type | *B* | *SE<sub>B</sub>* | t-value | *p* | RW | %*R*<sup>2</sup> |
| *R*<sup>2</sup> = .04 | VIQ | -0.13 | 0.01 | -10.16 | <.001 | 0.021 | 54.173 |
| *F* = 146.28, *p* < .05 | PIQ | -0.11 | 0.01 | -8.87 | <.001 | 0.018 | 45.827 |

*Note.* *B* = regression weight, *SE<sub>B</sub>* = standard error for *B*; RW = raw relative weight; %*R*<sup>2</sup> = Rescaled raw relative weight as a percent of total variance explained by model.

### Heterogeneity Analyses

In addition to calculating overall effect sizes for the associations between intelligence and antisocial variables, we also evaluated the heterogeneity, or between-studies variation, for each effect size (see Tables 1-4). Heterogeneity analyses revealed a high degree of between-study variation, suggesting larger differences than might be expected due to chance (*Q_B_*= 604.47, p= .000, I^2^ = 83%) among studies examining FSIQ and psychopathy. As a result, there may be moderators exerting an impact on study findings (a point we discuss in detail below). Similarly, effect sizes were heterogeneous for psychopathy with VIQ (*Q_B_*= 255.98, p= .000, I^2^ = 85%), and for PIQ (*Q_B_*= 77.01, p= .000, I^2^ = 65%). Further analyses revealed that the effect sizes were significantly heterogeneous for Factors 1 and 2 of psychopathy with intelligence and its facets. The I^2^ values ranged from 66% (for Factor 2 and FSIQ) to 94% (for Factor 2 and VIQ). Similarly, stratification by facets did not reduce the degree of heterogeneity (I^2^ ≥ 80%), except for effect sizes for facets 2 (I^2^ = 53%) and 3 (I^2^ = 62%) in relation to PIQ. Heterogeneity for Boldness/Fearless Dominance was moderate to high and was small to non-existent for Impulsive-Antisociality/Meanness and Disinhibition. For the remaining antisocial categories, analyses revealed that the effect sizes were significantly heterogeneous for FSIQ with ASPD (*Q*=50.57, p= .000; I^2^ = 74%) and CD (*Q*=458.00, p=.000; I^2^ = 95%). Heterogeneity analyses were not significant for ASPD and VIQ (*Q_B_*= .52, p= .914, I^2^ = 0%) or PIQ (*Q_B_*= 2.30, p= .313, I^2^ = 13%), but effect sizes were significantly heterogeneous for CD and both VIQ (*Q_B_*= 43.34, p= .000, I^2^ = 86%) and PIQ (*Q_B_*= 54.73, p= .000, I^2^ = 94%).

As alluded to, heterogeneous effect sizes can emerge for a couple of reasons, including simply by chance. However, differences in effect sizes might also represent the presence of moderating variables, or the fact that studies are not, in reality, measuring the same outcomes. Our analyses seemed to suggest that chance variation was unlikely to explain all of the heterogeneity observed. Therefore, we conducted moderator analyses for all effect sizes, with the exception of those related to ODD.

### Moderator Analyses

Moderator analyses were carried out in order to further evaluate which aspects of the studies included in our review (e.g., variation in measures used; age or gender of the sample) might be contributing to the variation we observed across studies. Results from categorical moderator analyses are presented in Table 6 for ASPD, CD and psychopathy. In total, we examined ten potential categorical moderators including category type (ASPD, CD, or psychopathy), antisocial spectrum disorder type (ASPD and CD), gender, age group, psychopathy or antisocial personality measure, IQ measure, recruitment site, covariates, region, and publication type) and one continuous variable (year of publication).

**Table 6.**
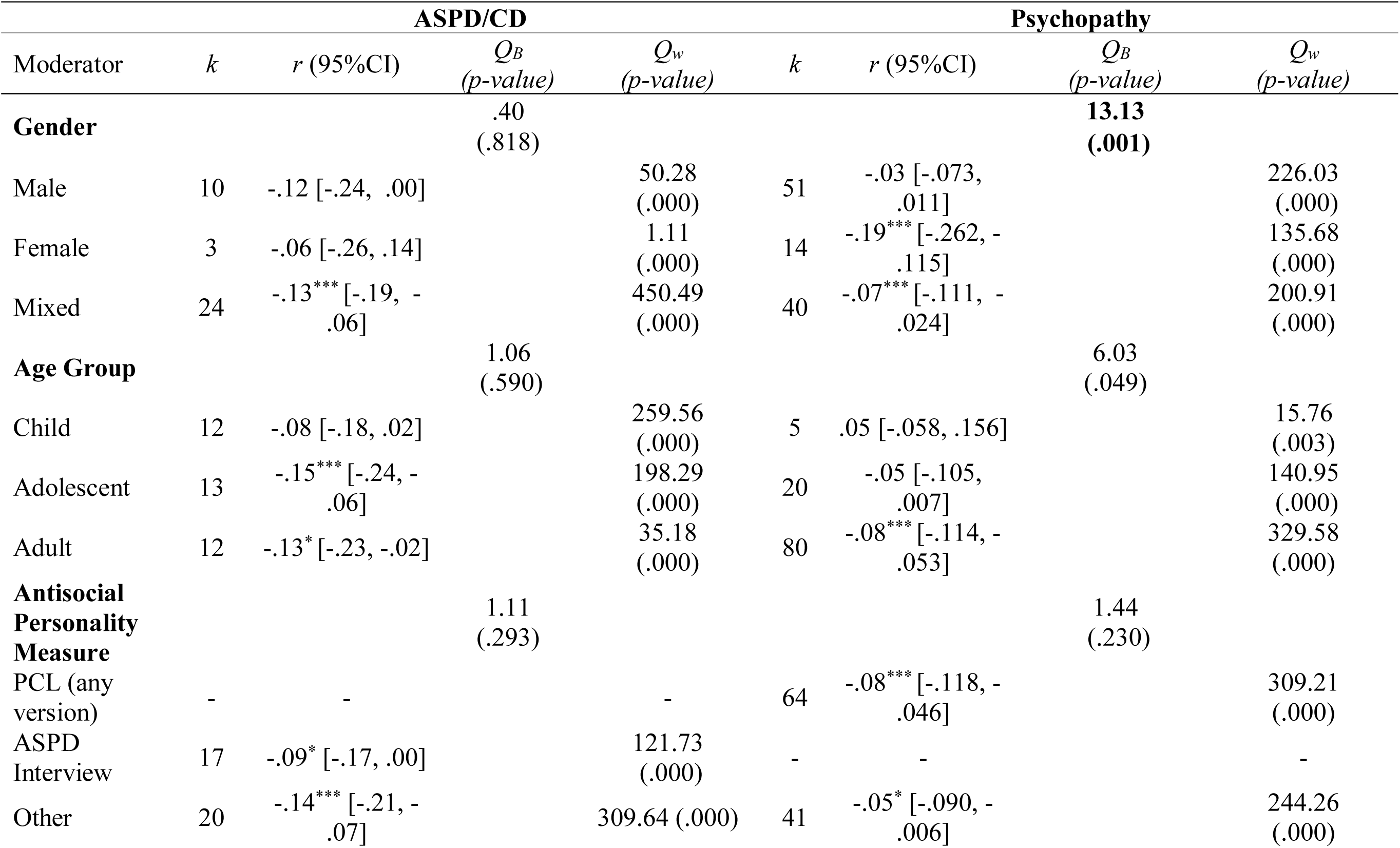
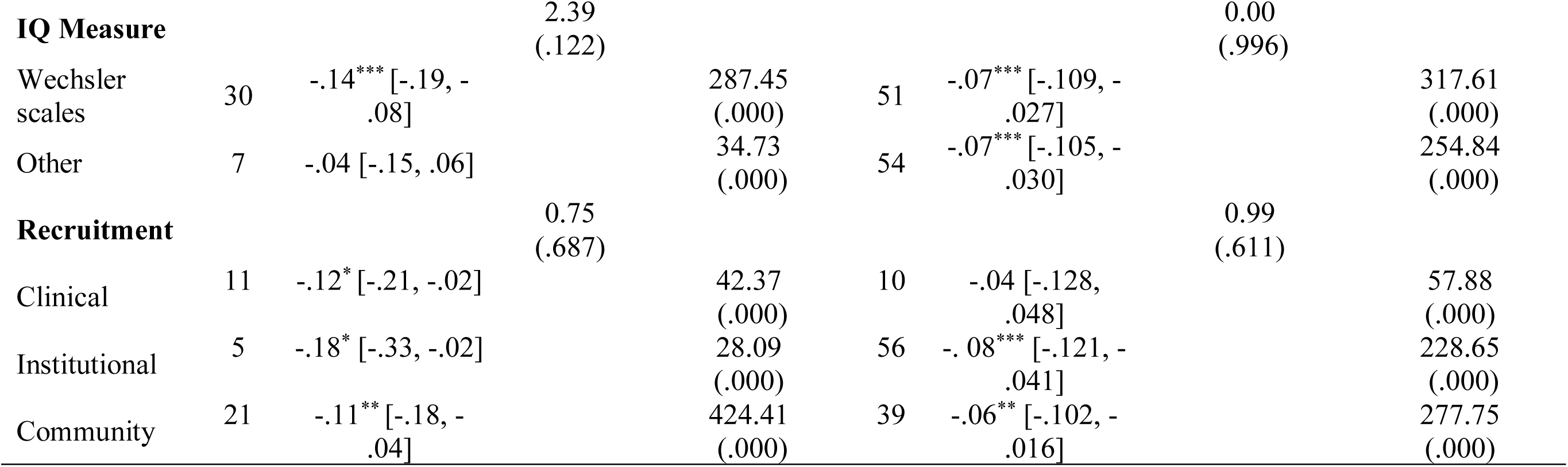
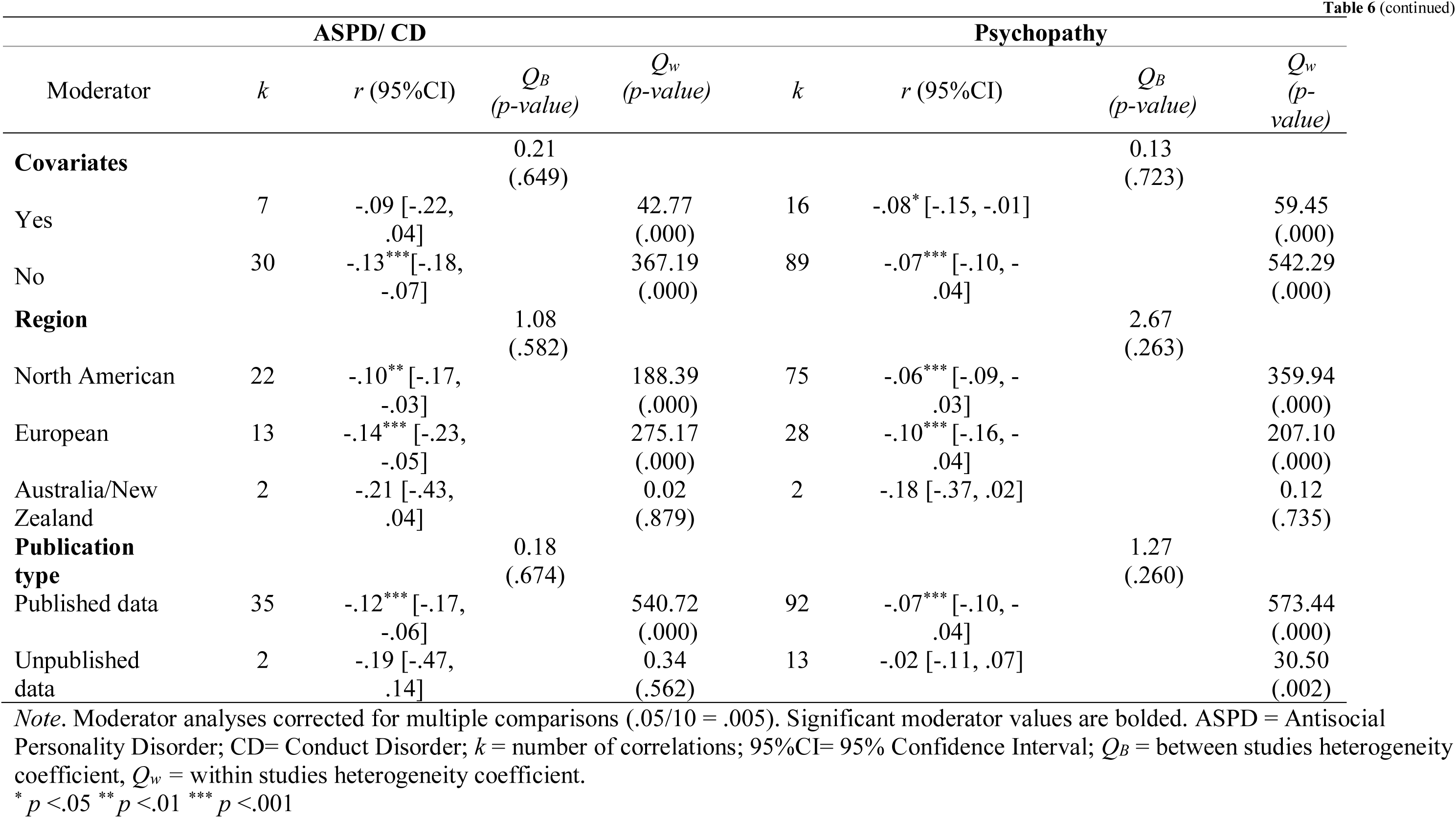
Results of categorical moderator analyses under random effects model for ASPD, CD, and psychopathy

Running multiple analyses on the same sample (here the collection of studies) increases the risk for Type I error (a false positive; Bender & Lange, 2001). Therefore, we corrected the significance level of our broad construct (ASPD/CD and psychopathy) moderator analyses for multiple comparisons (.05/10= .005). First, we conducted two analyses to test the possibility that the type of antisocial category moderated the results between intelligence and a given outcome (either psychopathy, ASPD, or CD). Put differently, this portion of the analysis examined whether the effects observed between intelligence, and each outcome, differed from one another. In the first analysis, we included all three categories ASPD, CD, and psychopathy, but found no evidence that the effect differed between the categories (*Q_B_ =* 2.73, p= 0.255). We also conducted a second test to further examine potential differences between the two more behaviorally based categories (ASPD and CD), but again found no evidence for moderation (*Q_B_* = 0.09, p= 0.771). Due to the conceptual overlap between CD and ASPD and the lack of significant heterogeneity between the two categories, the remainder of the moderator analyses were conducted on CD and ASPD together (see Table 6). None of the eight categorical moderators we tested (gender, age group, measure of antisocial traits, IQ measure, recruitment site, whether covariates were included, region, and publication type) emerged as statistically significant. Finally, meta-regression analysis revealed that year of publication was not associated with between-studies variation in effect sizes (B= 0.00, SE= 0.003, p= .142).

With regards to psychopathy, one categorical moderator variable was significant in our analysis of the relationship between psychopathic traits and FSIQ after correcting for multiple comparisons (.05/10 = .005). (see Table 6). There appeared to be a gender difference in the association between psychopathy and FSIQ, as gender was found to significantly moderate the association (*Q_B_*=13.13, p = .001). Specifically, female samples demonstrated significant small negative associations between psychopathy and FSIQ (r= -.19, p= .000), whereas no significant association was found for males (r= -.03, p= .145). Mixed samples yielded a significant, small effect size (r= -.07, p= .002). None of the other seven categorical moderators tested (age group, psychopathy measure, IQ measure, recruitment site, inclusion of covariates, region, and publication type) demonstrated a significant effect. Finally, meta-regression analysis suggested that the effect size was not significantly associated with the publication year (*B*= 0.00, SE= 0.001, p=.377). Unfortunately, there were not enough studies examining the association between psychopathy and VIQ or PIQ to allow for assessing moderators of those relationships.

We next conducted moderator analyses for the studies examining FSIQ, VIQ, and PIQ in relation to Factors 1 and 2 of psychopathy (see Table 7). In order to reduce the chance of Type 1 error, all moderator analyses were corrected for multiple comparisons (.05/8 = .006). The associations between FSIQ and Factors 1 and 2 of psychopathy were not moderated by any variable that we considered. The association between Factor 1 psychopathy and VIQ was not moderated by any of the eight moderators tested (i.e., gender, age group, psychopathy or measure, IQ measure, recruitment site, inclusion of covariates, region, and publication type). Conversely, the age of the sample (*Q_B_* = 18.379, p= .000) moderated the association between Factor 2 and VIQ. With regards to sample age, both adults (r= -.21, p= .000) and adolescents (r= -.20, p= .002) yielded significant (moderate) correlations, while samples consisting of children yielded no effect (r= .08, p= .172).

**Table 7.**
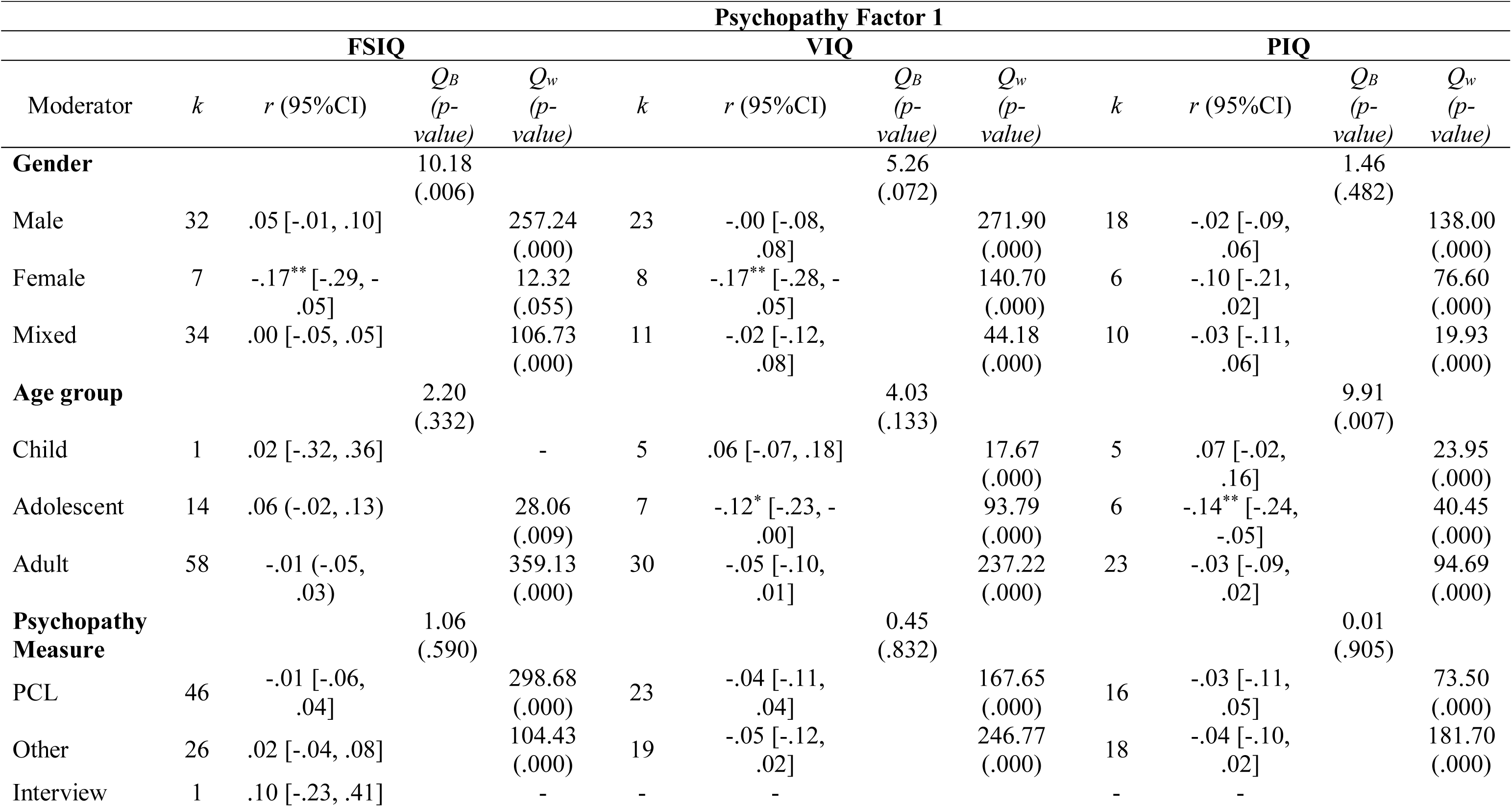
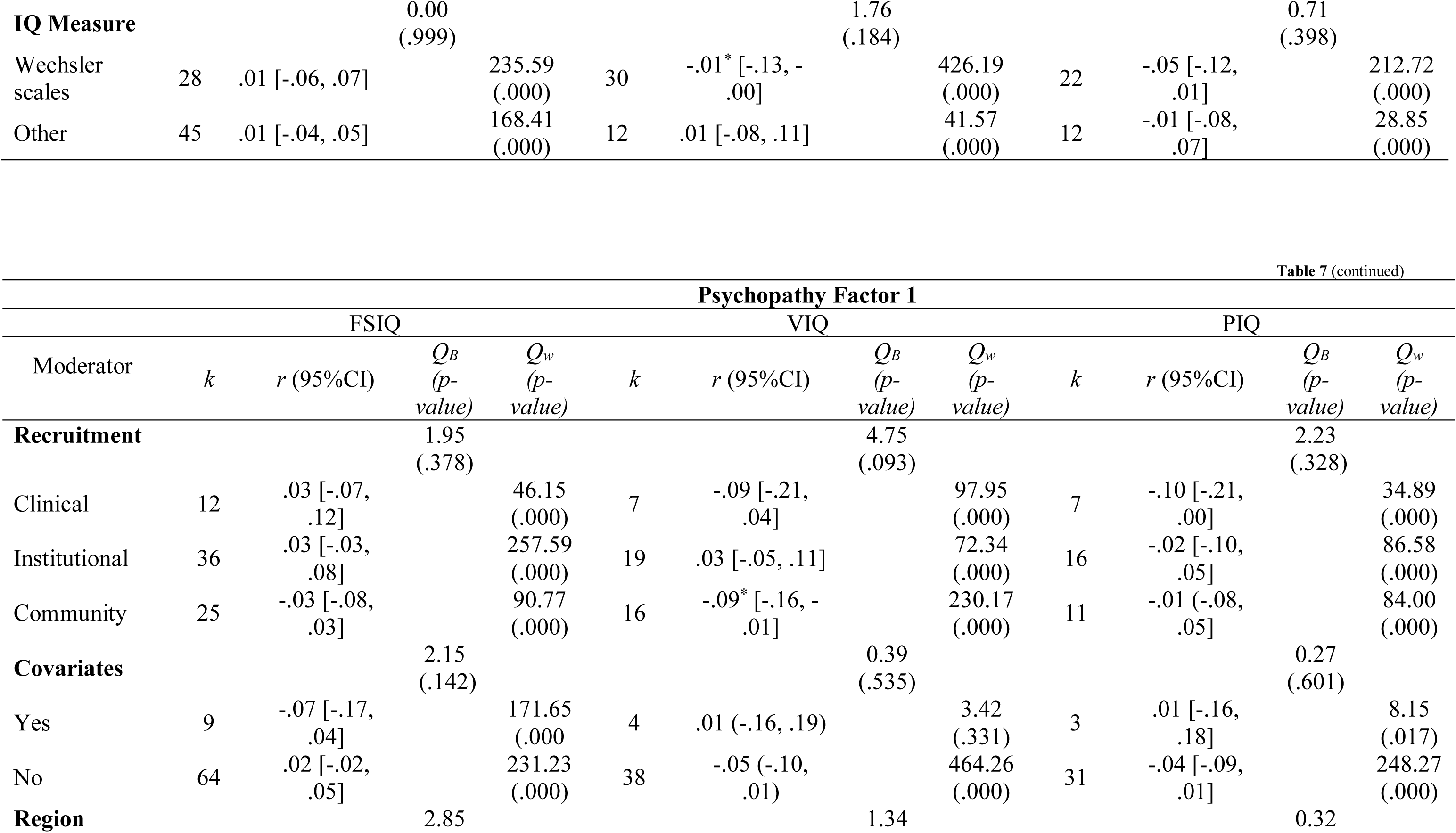
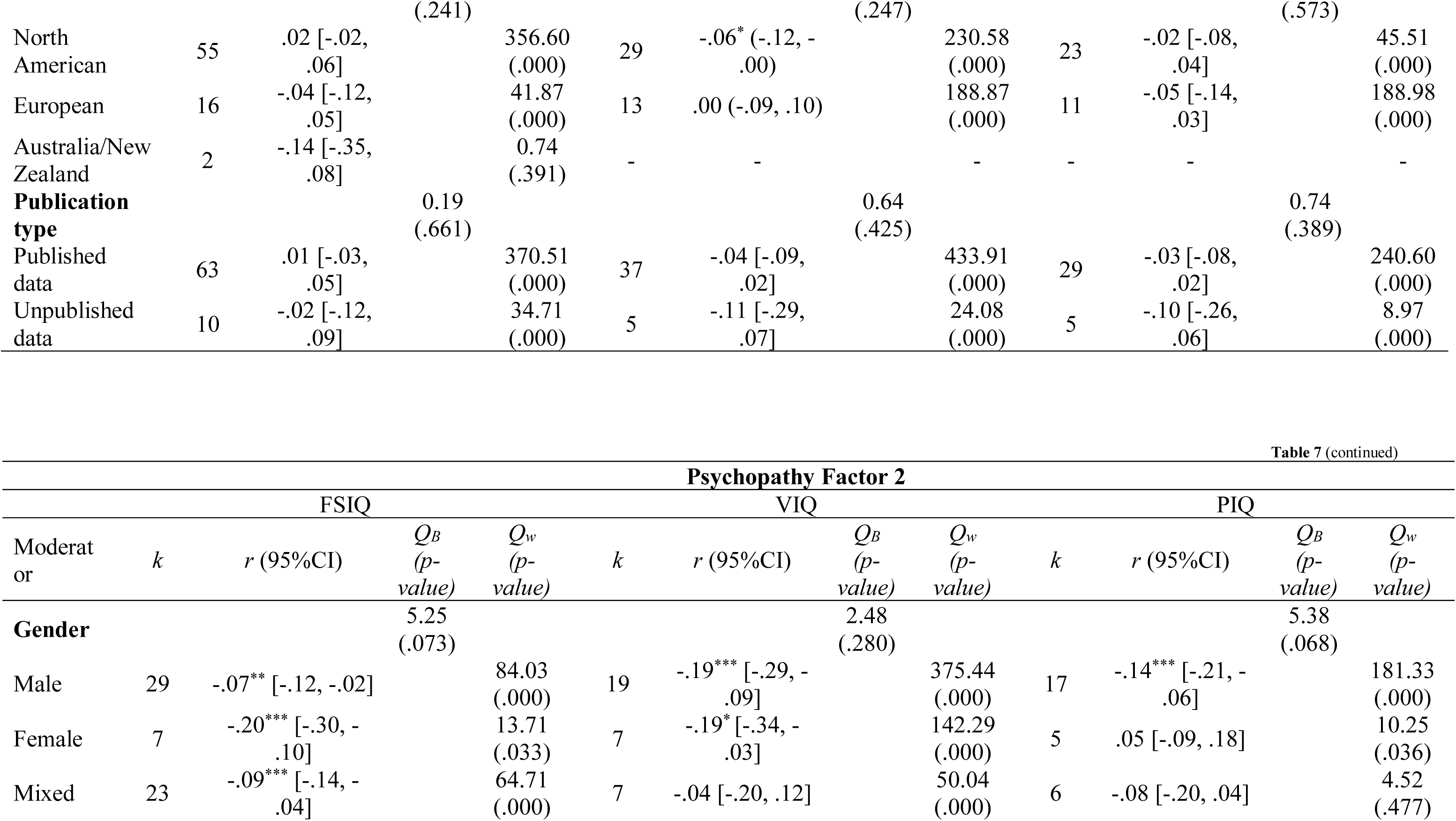
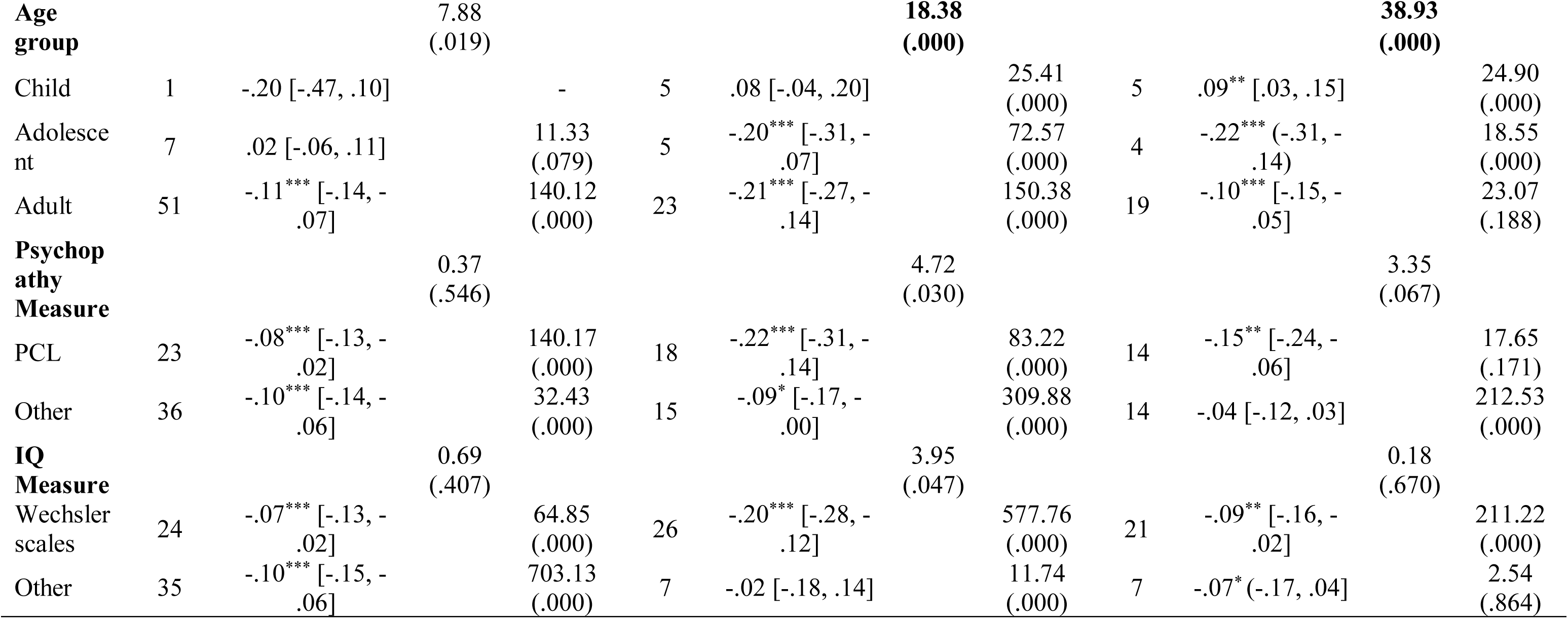
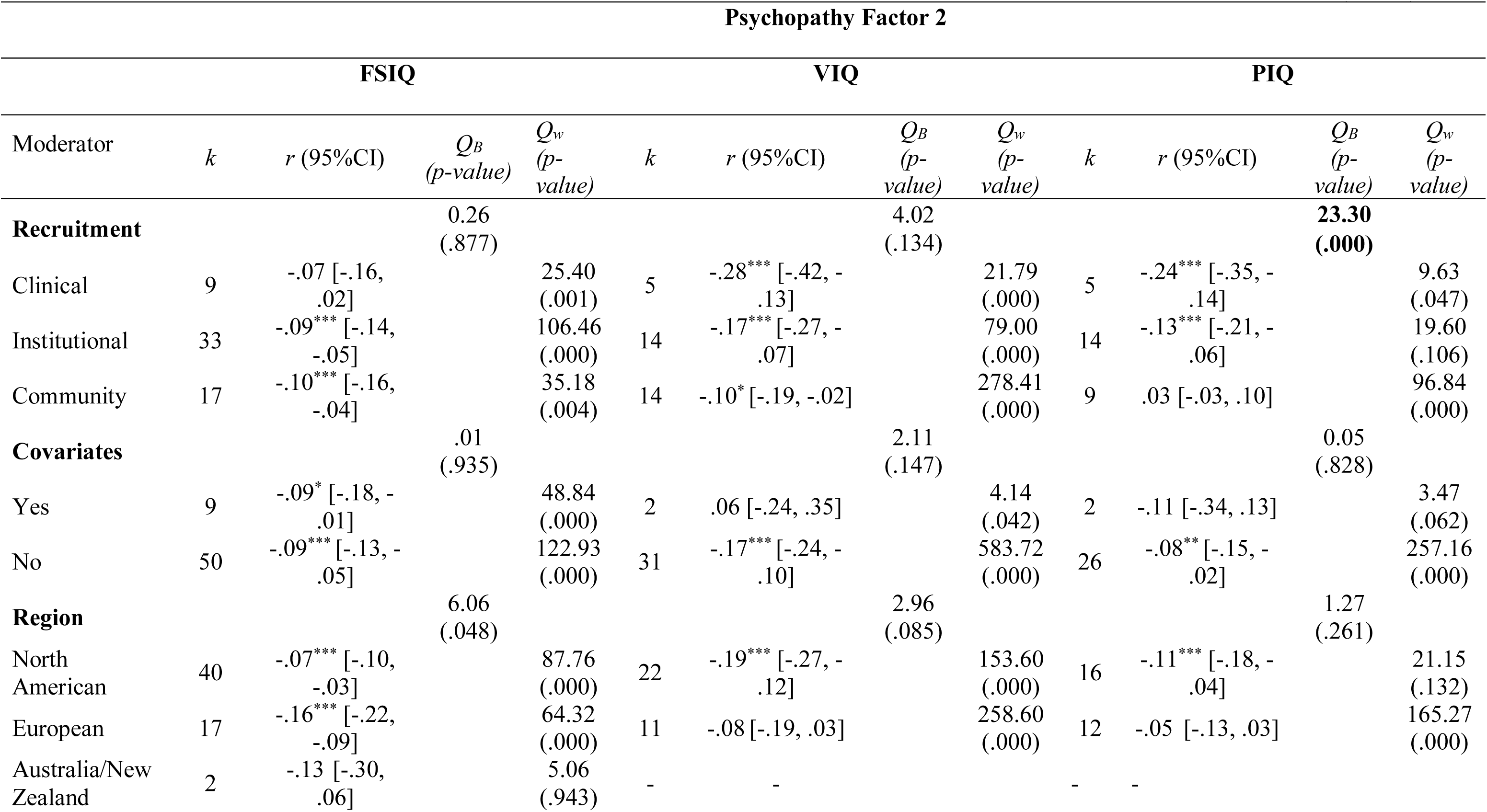
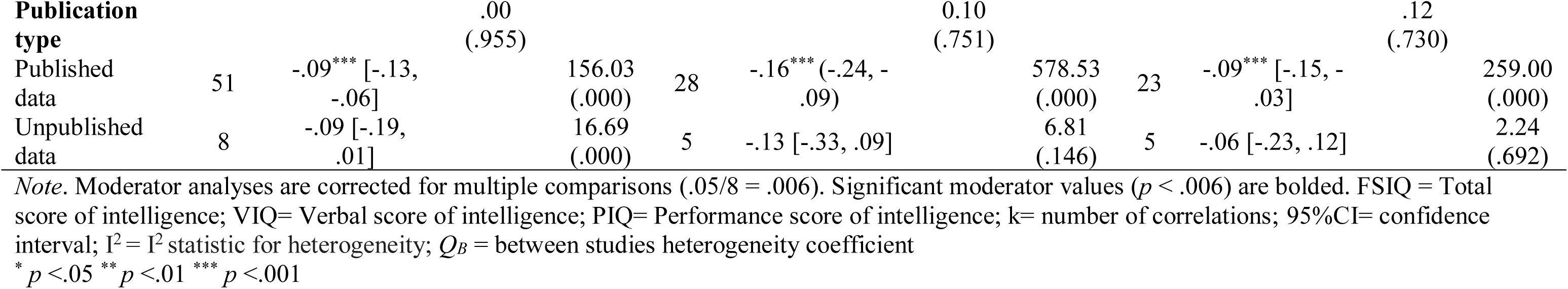
Results of categorical moderator analyses under the random effects model for Factor 1 and 2 of psychopathy and subtypes of intelligence

When examining effect sizes for the relationship between Factor 1 psychopathy and PIQ, no significant moderators were found. With regards to Factor 2 psychopathy and PIQ, the age of the sample (*Q_B_* = 38.93, p= .000) and recruitment site (*Q_B_* = 23.30, p= .000) were significant moderators. Adolescents yielded a moderate, negative effect size (r = -.22, p=.000) and adults yielded a small, negative effect size (r= -.10, p= .000), whereas children evinced a small, positive effect size (r = .09, p= .004). Second, a moderate effect size was found for the clinical sample (r = -.24, p= .000) and a weak effect was found for institutional samples (r = -.13, p= .001), whereas the community sample evinced no association (r = .03, p= .312). Meta-regression analyses revealed that the effect sizes for Factor 1 and Factor 2 with VIQ and PIQ were not significantly associated with the publication year (Factors 1 and 2, and VIQ: *B*= -.00, SE= .005, p= .715; *B*= -.01, SE= .007, p= .393, respectively. Factors 1 and 2, and PIQ: *B*= .01, SE= .005, p= .913; *B*= -.01, SE= .005, p= .243, respectively). Due to the small number of effect sizes available for analysis, we could not conduct moderator analyses for facets of psychopathy.

### Publication Bias and Sensitivity analyses

In order to examine the robustness of the results obtained in the current study, we examined the potential influence of publication bias using “Trim and Fill” analysis (Duval & Tweedy, 2000) and Eggeŕs regression test. Briefly, the “Trim and Fill” technique utilizes a funnel plot as a method of gauging the asymmetry in published work. Funnel plots are particularly useful as they provide a scatter plot of the various effect sizes from studies included in a meta-analysis. If a body of research is generally free from publication bias, we might expect that the effect sizes reported by the various studies would be normally distributed around the overall (average) effect size. Any asymmetry in the scatter plot, on the other hand, is suggestive of publication bias.

Once the degree of asymmetry is assessed, the “Trim and Fill” technique is used to iteratively “trim” the most extreme small studies from the positive side until the plot becomes symmetrical around a new, adjusted effect size. Trimmed studies, then, are added back algorithmically, along with mirror images of the trimmed studies which are imputed on the opposite side (thus, the “fill” aspect of the procedure; see Duval & Tweedy, 2000) to visually represent the approximate values of the effect sizes missing due to publication bias (Duval & Tweedie, 1998; 2000). Egger’s regression test regresses an estimate’s standard normal deviate (SND; the odds ratio divided by the standard error) on to its precision (the inverse of the standard error) so as to provide an estimate of bias in the funnel plot (see Egger et al., 1997).

First, the random effects “Trim and Fill” analysis for the other antisocial disorders (ASPD and CD combined), determined that zero studies had to be added on either the right or the left side to be symmetrical (see Figure 2A). However, the Eggeŕs regression test failed to rule out the possibility of heterogeneity, perhaps owing to publication bias (*B*= −2.76, SE= 0.869, *p*= .003). Next, the “Trim and Fill” analysis imputed zero studies on either side for psychopathy and intelligence (see Figure 2B), and the Eggeŕs regression test for psychopathy did not report a significant slope coefficient (*B*= −0.47, SE= 0.434, *p*= .277). In other words, the “Trim and Fill” did not indicate publication bias is a major concern. In sum, evidence from the tests for publication bias suggest there may be some bias in the literature with regards to studies on intelligence and antisocial disorders/traits, but there does not seem to be evidence of publication bias for studies on intelligence and psychopathy.

**Figure 2:**
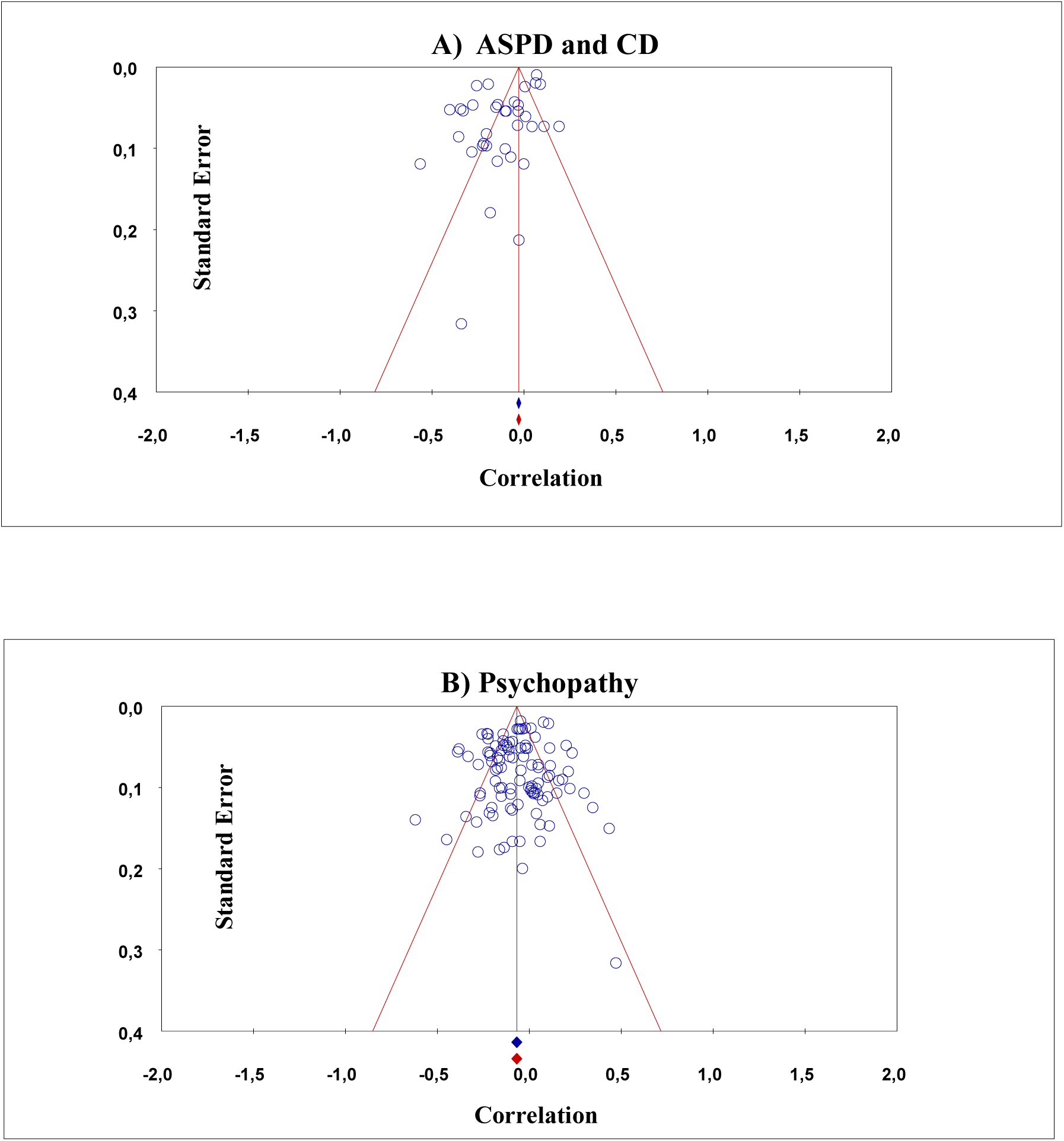
Funnel plot with trim-and-fill imputations for Full-scale intelligence and antisocial disorders combined (A), and for psychopathy (B).

At this point, we should note that all methods of estimating various aspects of publication bias have their own strengths and limitations. Along these lines, we also considered including p-curve and p-uniform analyses. While certainly useful in some scenarios when conducting meta-analyses, recent research has argued that in meta-analyses evincing high levels of heterogeneity, such as in the current study, these methods may not represent the most ideal approach. For example, a recent study by van Aert and colleagues (2016) conducted a series of simulations in order to examine if, and to what extent, p-curve and p-uniform analyses were vulnerable to varying degrees of heterogeneity in a set of results. In particular, van Aert et al. (2016: p.718) conclude that:

> “If the main goal of the meta-analysis is to estimate the average true effect of the whole population of studies in the presence of heterogeneity (I^2^ ≥ .5), we do not recommend using p-curve or p-uniform because doing so generally overestimates average true effect size (Recommendation 5a).”

Given the degree of heterogeneity in our own study, we opted against employing either the p-curve or p-uniform analyses.

As a second test of robustness, however, we also evaluated the impact of potential influence of outlier studies. First, we assessed the effect of four studies (Ford, et al. 2007; Hofvander et al. 2011; Masten et al. 1999; and Nomura et al., 2008) that used antisocial behavior questionnaires that were not associated with a specific psychiatric diagnosis. These studies were included because the items were similar to those associated with diagnostic criteria and removing them from our analyses had no substantive effect on the results.

When examining extreme values, we found one study to be an outlier for psychopathy and intelligence. Specifically, Nestor et al. (2005) reported the strongest positive correlation between psychopathy and intelligence and a high standard error (r= .47, SE= .32 N= 13). For ASPD, we similarly found one study, Pera-Guardiola et al. (2016) that reported a negative effect size that was fairly strong with an accompanying high standard error (r= -.34, SE= .32, N=13). Nonetheless, the exclusion of these outliers did not have an effect on the total effect size (r= -.08, p= .001), nor did it have an impact on the antisocial sub-types effect sizes (psychopathy: r= -.07, p= .001; ASPD: r= -.13, p= .001), which is likely due to the small sample sizes of the outlier studies.

## Discussion

### Intelligence and Psychopathy

The current meta-analysis presents the most comprehensive statistical evaluation to date of the association between intelligence and psychopathy, including analysis by factor and facet, as well as between intelligence and various antisocial disorders. Despite previous meta-analytic work suggesting that no relationship exists between intelligence and psychopathy (O’Boyle et al., 2013), our analyses suggested that a small, yet statistically significant, negative relationship exists between indicators of intelligence and psychopathy broadly conceptualized (*r* = -.07). Finer grained analysis of intelligence through indicators of verbal and performance IQ, revealed that psychopathy was significantly and negatively associated with both VIQ and PIQ; however, results of MASEM analyses suggested that VIQ is the primary aspect of intelligence associated with psychopathic traits.

Arguably of greater importance, owing to the growing recognition of the multi-dimensional nature of psychopathy (see Lilienfeld, 2018), we sought to examine if, and to what extent, aspects of psychopathy were associated with intelligence. The interpersonal and affective (Factor 1) aspects of psychopathy were statistically unrelated to intelligence (FSIQ, VIQ, & PIQ), but the antisocial and impulsive Factor 2 was associated with lower scores on FSIQ and both of its facets. These findings suggest that lower intelligence is particularly associated with the behavioral problems observed in those who score higher on measures of psychopathy or are diagnosed with antisocial disorders, and is in line with findings suggesting that intelligence is negatively related to criminal offending (Schwartz et al., 2015), aggression (Kennedy, Burnett, & Edmonds, 2011), and impulsivity (Lynam, Moffitt, & Stouthamer-Loeber, 1993; Meldrum et al., 2017).

Investigation of factors of psychopathy still only provides a coarse level of analysis, so we next examined associations between intelligence and the four-factor structure of psychopathy. Associations with the interpersonal facet showed a significant, *positive*, small effect size for FSIQ. There was not a statistically significant association with either VIQ or PIQ, although this may be due to the small number of effect sizes. The affective facet was weakly and negatively related to all aspects of intelligence considered. Finally, both the lifestyle and antisocial facets were negatively related to both VIQ and PIQ, but neither was statistically related to FSIQ. The lack of a significant association between these two facets and FSIQ is surprising as one might assume that there would be a significant correlation to match the correlations at the domain level of intelligence (i.e., VIQ and PIQ). Furthermore, the lifestyle and antisocial facets are most closely related to antisocial behavior, which we noted earlier is consistently related to intelligence in a negative direction. Thus, it seems likely that the lack of a significant association found here between FSIQ and these two facets of psychopathy may be due to a statistical artifact or some (unknown) moderating effect that we are unable to test here. The consistent negative associations between VIQ and higher scores on the affective, lifestyle, and antisocial facets of psychopathy, combined with the results of the MASEM analyses which attributed most of the variance in psychopathy accounted for by intelligence to VIQ, is not particularly surprising as it dovetails with previous suggestions (e.g., DeLisi et al., 2010) that many aspects of psychopathy are actually related to lower verbal ability, in contrast to the descriptions of Cleckley (1941).

The association between intelligence and the interpersonal facet was positive, indicating that those who are more superficially charming and manipulative may actually demonstrate somewhat higher levels of intelligence. This is not to suggest that superficially charming and manipulative personality types are among the highest scoring individuals on measures of intelligence, only that they do not seem to possess any severe cognitive deficits. To the extent that these findings continue to replicate, it also seems plausible that it was this aspect of psychopathy, which led Cleckley (1941) to his original description of “psychopaths” as possessing “good intelligence.” It is possible that such an association exists, despite our finding of an overall negative association of global psychopathy with intelligence, which does not lend support his description.

Finally, with regards to aspects of psychopathy and intelligence, we examined conceptualizations that include a boldness or fearless dominance component. As noted previously, the inclusion of boldness/fearless dominance as a psychopathic trait has been intensely debated (e.g., Lilienfeld et al., 2012; Miller & Lynam, 2012), and unfortunately the current study cannot resolve the debate. Yet, in light of a prior meta-analysis finding suggesting that boldness correlated moderately with many other psychopathy measures (Lilienfeld et al., 2016), we felt it was relevant to also examine the associations between intelligence and conceptualizations of psychopathy that include a boldness/fearless dominance component. To do so, we combined the Fearless Dominance facet of the PPI and the Boldness domain of the triarchic model (effect sizes were only available for studies using triarchic scales created within the MPQ) and found a modest, positive association with FSIQ. We did not find a statistically significant association with either VIQ or PIQ. Conversely, impulsive-antisociality from the PPI and triarchic meanness and disinhibition (MPQ-Tri) subscales were combined and demonstrated a weak, negative relationship with all types of intelligence. These results add an additional layer of evidence for the multi-dimensionality of psychopathy and for the interpretation of boldness/fearless dominance as related to adaptive functioning (see Lilienfeld et al., 2016).

### Intelligence and other Antisocial Constructs

Because psychopathy is not officially recognized in the DSM but is closely related to and often treated as nearly (or completely) interchangeable with several DSM constructs (ASPD, CD, and ODD), we also examined the associations between intelligence and these antisocial disorders. Our results indicated that FSIQ was negatively associated with ASPD and CD, but positively with ODD (although we are hesitant to interpret the effect sizes for ODD and many of the analyses for VIQ/PIQ with CD and ASPD due to the small number of effect sizes available for analysis). Unfortunately, we were unable to parse shared and unique associations between psychopathy, ASPD, and CD with intelligence (the way we assessed VIQ and PIQ with each of these constructs) because we did not have the requisite meta-analytic associations between them (it is also important to note that interpreting the results of such an analysis would be difficult due to the extent to which partialling alters a construct; see Lynam et al., 2006).

We did, however, examine the confidence intervals for the overall effect sizes for the association between intelligence and each construct to assess if the associations appeared to differ in magnitude, but there does not appear to be a significant difference as the confidence intervals are overlapping. Thus, we did not find evidence for differential associations between intelligence and the antisocial constructs we considered (with the exception of ODD), although it is important to note the evidence we found for differential associations between intelligence and facets of psychopathy. Therefore, analyses that compare global psychopathy with these related constructs may be missing critical information should they exclude an analysis of the facet level measurements.

### Heterogeneity and Moderator Analyses

Importantly, our results also uncovered a moderate to high degree of heterogeneity between studies, suggesting that moderating variables may explain differences across studies. Indeed, moderator analyses revealed that the type of sample (community, institutional, or clinical), age, and gender were conditioning several of the associations being tested, some in unexpected ways. Given the unexpected nature and lack of consistency of moderating effects across various aspects of psychopathy and intelligence, we want to emphasize that our interpretations are highly speculative, and we urge awaiting future research assessing the replicability of these results before stronger conclusions are drawn. With regards to psychopathy total scores, moderator analyses suggested that the gender of the samples affected the strength of the relationship between FSIQ and psychopathy. Somewhat surprisingly, female samples demonstrated a moderate negative relationship between intelligence and psychopathy total scores, while male samples demonstrated no overall relationship and mixed samples produced a weak, negative relationship. Furthermore, examination of the confidence intervals for the male and female samples indicated they were not overlapping, and thus were significantly different.

The difference that emerged between male and female samples was interesting and is something that merits some discussion (as well as additional research). On one hand, males may simply demonstrate less variation in and score lower on certain traits, such as empathy (relative to females; see Baron-Cohen, 2002; Baron-Cohen & Hammer, 1997). A lack of empathy, in particular, is a key aspect of psychopathic tendencies, thus potentially resulting in less male variability on measures of psychopathy. The consequence of this, moreover, is that it could become more difficult to detect what is already a relatively small association with intelligence. Another potential explanation is that there is sampling bias. Institutional samples, especially samples of incarcerated males, are overrepresented in this literature. Incarcerated samples tend to demonstrate higher levels of psychopathic traits compared to other types of samples, such as community samples (e.g., Coid, et al., 2009). It is a possibility that the abundance of incarcerated male samples, either independent of, or jointly with reduced male variability on psychopathic traits, biased the meta-analytic effect size through an effect similar to range restriction. However, it must be noted that moderator analyses did not indicate a significant difference between institutional and other sample types for most effect sizes (though the test for moderation by sample type does not control for gender). Furthermore, some evidence suggests males tend to be *more* variable with regards to intelligence compared to females (e.g., Johnson et al., 2008; 2009). Greater variability in intelligence in a pooled sample that is potentially less variable on psychopathic traits and skewed towards the higher end of the distribution for that construct could all be contributing to the lack of an association for males.

The above is admittedly speculative, however, and at this point further speculation seems unhelpful. More fruitful insight will come from additional research examining various measures of cognitive ability and psychopathic tendencies in both males and females separately. In particular, research will be needed which examines potential gender differences in the association between intelligence and psychopathy at the facet level, given our findings that total psychopathy scores are obscuring important variability.

We also found an interesting moderating effect of age on the associations between indictors of intelligence (VIQ and PIQ) and Factor 2 psychopathy, such that intelligence facets were inversely related to Factor 2 in adolescent and adult samples, but were statistically unrelated in samples of children (or positively related in the case of PIQ). One potential explanation of this moderating effect is related to the fact that Factor 2 is closely tied to antisocial behavior, which typically emerges most strongly in adolescence and extends into early adulthood (e.g., the age-crime curve, see Farrington, 1986). Therefore, it may be that samples of children simply do not display enough magnitude and variance in antisocial behavior (relative to adolescents and adults) for a statistical association to emerge. The finding of a positive association between PIQ and Factor 2 for children is a greater puzzle and unintuitive and we have decided not to speculate on its interpretation until additional research replicates the finding.

Finally, we also found that the type of sample (clinical, institutional, or community) moderated the association between PIQ and Factor 2, with clinical and institutional samples evincing negative associations and community samples demonstrating a nonsignificant association. Assuming the effect is a true effect, we believe the same interpretation we applied in terms of the moderating effect of age could apply here. That is, clinical and institutional samples likely contain much greater variability in Factor 2 traits and antisocial behavior which may allow for a statistically significant association with intelligence to emerge. It is unclear, however, why the sample type would only moderate the association between PIQ and Factor 2 and not VIQ or FSIQ.

### Publication Bias and Sensitivity analyses

Publication bias and extreme values, among other factors, are always a concern in meta-analytical work due to their effects on estimated effect sizes. Therefore, we conducted multiple tests to assess for the possibility that these factors were biasing our results. Two measures of publication bias (“Trim and Fill” analysis, Duval & Tweedy, 2000; and Eggeŕs regression test) did not suggest any evidence of publication bias for psychopathy, but Egger’s regression failed to rule out bias for the related antisocial constructs. There are other methods of assessing bias, each with its own limitations, so conclusions asserting that there is no bias present in the intelligence and psychopathy literature should be avoided. Nonetheless, the current study found no evidence to suggest that publication bias is having a large impact. We also re-ran analyses with extreme studies removed to assess for outlier effects, but the results were not substantively altered.

### Limitations and Future Directions

Prior to concluding, there are some important limitations in the current study that will require attention in future research. We must reiterate that all non-Wechsler intelligence measures were collapsed together into a very broad “other” category. While this was done in order to streamline an already hulking analysis, it is admittedly not ideal given that these measures tap in to slightly different aspects of general intelligence. At the same time, it can be reasonably assumed that each measure loads on the same underlying construct (in varying degrees of magnitude) and are thus all capturing aspects of the same trait (Ritchie, 2015). Yet, collapsing them as we did is *not* the same thing as creating a global construct of general intelligence. It is entirely possible, then, that effects may vary from measure to measure, and testing whether this is the case is a question that remains in need of addressing. Additional work will be needed to further dissect whether, and to what extent, effects of non-Wechsler based tests differ when predicting the outcomes tested herein.

Related to this point, we did not include studies that used additional surrogates for intelligence testing, such as measures of working memory. Such measures are correlated with IQ and a strong argument could be made for their inclusion in the current meta-analysis. However, despite self-imposing a limitation on our study by including only standardized intelligence tests and subtests, we felt the tradeoff was necessary given the already large amount of material contained in this meta-analysis. Therefore, we opted to retain our current approach, and encourage further work examining additional measures of intelligence.

As a final point related to our treatment of intelligence in the current study, we must note that parsing intelligence into VIQ and PIQ is not intended to represent the standard research practices common in modern cognitive and intelligence literatures. Although early iterations of the Wechsler intelligence scales used the FSIQ, VIQ, and PIQ structure, more recent iterations have exchanged the VIQ and PIQ denotation in favor of a four-factor structure (verbal comprehension, working memory, processing speed, and perceptual reasoning; Wechsler, 2008). Additionally, researchers have also conceptualized intelligence in terms of “crystallized” and “fluid” intelligence (Cattell & Horn, 1978). Crystallized intelligence can be thought of as the accumulated knowledge of an individual, whereas fluid intelligence is comprised of analytical ability and problem solving (Cattell & Horn, 1978). To more easily summarize this body of work, we used the VIQ and PIQ distinction because many the studies found in our literature search used Wechsler instruments, especially older iterations of them. Thus, it is likely that future reviews of this literature will adequately make the distinction between fluid and crystalized intelligence in accordance with the changing paradigm around intelligence. Nevertheless, VIQ as used in the current paper can potentially be thought of as a rough, imperfect analogue to crystallized intelligence as it includes primarily tests of vocabulary and scholastically-based knowledge. Similarly, PIQ in the current study includes scores from tests such as Matrix Reasoning on the WAIS, which loads on a fluid intelligence factor (van Aken et al., 2017).

A second limitation of the current study relates to interpretations of causality. Overall, the existence of a negative relationship between these constructs may tempt the assumption that lower intelligence *causes* individuals to evince more psychopathic traits (or perhaps vice versa). While this is not beyond the realm of possibility, it is important to remember that we are only examining correlational data, and causal inferences must be avoided. Previous researchers, moreover, have suggested that intelligence might act as a mediator between psychopathic traits and antisocial behavior (Muñoz et al., 2008; Salekin et al., 2010). Kandel et al. (1988) found that higher intelligence, additionally, acts a protective factor against offending generally (i.e. not looking specifically at psychopathic individuals). Conversely, Muñoz et al. (2008) found higher intelligence to be a risk factor for increased violent offending among psychopathic individuals. Yet, Salekin and colleagues (2010) found no relationship between IQ and offending among those scoring highly on psychopathy.

It is also plausible that if a causal connection exists, higher levels of psychopathic traits may ultimately serve as a barrier to environmental exposures that could increase levels of intelligence. For example, if psychopathic individuals miss more school due to truancy, suspension, or incarceration, or ultimately complete fewer years of education, then they may fail to reap the full intelligence boosting effects of education (Ritchie & Tucker-Drob, 2018). Importantly, a similar scenario might be posited for intelligence and ASPD and conduct disorder. Genetic confounding, too, might play an important role, such that pleiotropic genetic influences might both increase psychopathic tendencies, while also lowering cognitive ability (and a similar possibility exists for intelligence and antisocial disorders) (Barnes et al., 2014). However, genetically sensitive designs will be required to further examine this interesting possibility, and more work is needed in general to fully unpack causal pathways (Barnes et al., 2014). Although, evidence shows that variation across both intelligence and psychopathy measures are impacted by genetic variation (Deary, 2013; Ferguson, 2010; Gunter, Vaughn & Philibert, 2010; Plomin & Deary, 2015), it remains a challenge to identify the (numerous) polymorphisms responsible for the heritability of psychopathy (Viding et al. 2013) and their association with the many alleles that might influence intelligence (Lee et al., 2018).

## Conclusion

Overall, we found small, negative associations between intelligence and psychopathy, as well as between intelligence and antisocial disorders and conduct disorder (with the exception of ODD). These associations did not appear to differ in magnitude, suggesting that intelligence does not differentiate these constructs, at least at the global level. Importantly, these relationships were significantly, albeit inconsistently, moderated by certain variables, such as age and gender, and differed across the facets of each construct being examined. The association for global psychopathy and FSIQ was negative for both males and females, but was greater in magnitude for females; however, gender did not moderate associations with VIQ or PIQ, Factor 1 or 2 of psychopathy, or any of the other antisocial constructs. Similarly, age of sample (children, adolescents, or adults) was found to moderate the association of VIQ and PIQ, but not FSIQ, with Factor 2 psychopathy, but not Factor 1 or psychopathy total scores. Facet level examination of psychopathy provided strong evidence in support of the multi-dimensional nature of psychopathy and differential correlates between the dimensions. An examination of psychopathy conceptualizations that include a boldness/fearless dominance component also produced evidence in support of the potentially adaptive nature of the boldness/fearless dominance component, as it was positively associated with intelligence. Future research should seek to further unpack the neurobiological, genetic, and evolutionary underpinnings and covariation among these constructs at the facet level, as well as the moderators that affect them. In particular, longitudinal and genetically sensitive designs are needed to bring us closer to causal inference and understanding how these constructs might interact across development and the life-course. For now, what remains clear is that as overall psychopathy scores increase, intelligence, generally speaking, does not.

References marked with an asterisk indicate studies included in the meta-analysis.

## Footnotes

1 From: Moher D, Liberati A, Tetzlaff J, Altman DG, The PRISMA Group (2009). Preferred Reporting Items for Systematic Reviews and Meta-Analyses: The PRISMA Statement. PLoS Med 6(7): e1000097. doi:10.1371/journal.pmed1000097. For more information, visit www.prisma-statement.org.

## References

*Allen, J. L., Briskman, J., Humayun, S., Dadds, M. R., Scott, S. (2013). Heartless and cunning? Intelligence in adolescents with antisocial behavior and psychopathic traits. Psychiatry Research, 210(3), 1147–1153.

American Psychiatric Association. (1987, 1994, 2000, 2013). Diagnostic and statistical manual of mental disorders-text revision (3rd ed., revised ed.; 4th ed.; 4th ed., revised version; 5th edition). Washington, DC: APA.

Anderson, J. L., Sellbom, M., Wygant, D. B., Salekin, R. T., & Krueger, R. F. (2014). Examining the associations between DSM-5 Section III antisocial personality disorder traits and psychopathy in community and university samples. Journal of Personality Disorders, 28(5), 675–697.

*Andrade, J.T. (2009). Psychosocial precursors of psychopathy in a psychiatric sample: A structural equation model analysis. PhD Disseration, Boston College.

*Anton, M.E., Baskin-Sommers, A.R., Vitale, J.E., Curtin, J.J., & Newman, J.P. (2012). Differential effects of psychopathy and antisocial personality disorder symptoms on cognitive and fear processing in female offenders. *Cognitive*, Affective and Behavioral Neuroscience, 12(4):761–76. DOI: 10.3758/s13415-012-0114-x.

*Arseneault, L., Kim-Cohen, J., Taylor, A., Caspi, A., & Moffitt, T.E. (2005). Psychometric evaluation of 5- and 7-year-old children’s self-reports of conduct problems. Journal of Abnormal Child Psychology, 33(5), 537–550. DOI: 10.1007/s10802-005-6736-5.

*Bagshaw, R., Gray, N.S., Snowden, R. J. (2014). Executive function in psychopathy: The Tower of London, Brixton Spatial Anticipation and the Hayling Sentence Completion Tests. Psychiatry Research, 220(1), 483–489.

Barnes, J. C., Boutwell, B. B., Beaver, K. M., Gibson, C. L., & Wright, J. P. (2014). On the consequences of ignoring genetic influences in criminological research. Journal of Criminal Justice, 42(6), 471–482.

Baron-Cohen, S. (2002). The extreme male brain theory of autism. Trends in cognitive sciences, 6(6), 248–254.

Barry, C. T., Frick, P. J., DeShazo, T. M., McCoy, M., Ellis, M., & Loney, B. R. (2000). The importance of callous–unemotional traits for extending the concept of psychopathy to children. Journal of abnormal psychology, 109(2), 335–340.

Bartels, M., van Weegen, F. I., van Beijsterveldt, C. E., Carlier, M., Polderman, T. J., Hoekstra, R. A., & Boomsma, D. I. (2012). The five factor model of personality and intelligence: A twin study on the relationship between the two constructs. Personality and Individual Differences, 53(4), 368–373.

*Baskin-Sommers, A.R., Brazil, I.A., Ryan, J., Kohlenberg, N.J., Neumann, C.S., & Newman, J.P. (2015). Mapping the association of global executive functioning onto diverse measures of psychopathic traits. Personality Disorder, 6(4), 336–346. DOI: 10.1037/per0000125.

*Bate, C., Boduszek, D., Dhingra, K., & Bale, C. (2014). Psychopathy, intelligence and emotional responding in a non-forensic sample: an experimental investigation. The Journal of Forensic Psychiatry & Psychology, 25(5), 600–612.

*Beggs, S.M., & Grace, R.C. (2008) Psychopathy, intelligence, and recidivism in child molesters: Evidence of an interaction effect. Criminal Justice Behavior, 35(6), 683–695 683. DOI: 10.1177/0093854808314786.

Bender, R., & Lange, S. (2001). Adjusting for multiple testing—when and how?. Journal of clinical epidemiology, 54(4), 343–349.

*Benning, S. D., Patrick, C. J., Blonigen, D. M., Hicks, B. M., & Iacono, W. G. (2005). Estimating facets of psychopathy from normal personality traits: a step toward community. *Epidemiological Investigations*. Assessment, 12(1), 3–18.

*Benning, S. D., Patrick, C.J., Hicks, B.M., Blonigen, D.M., Krueger, R.F. (2003). Factor structure of the psychopathic personality inventory: validity and implications for clinical assessment. Psychological Assessment, 15(3), 340–350. DOI: 10.1037/1040-3590.15.3.340.

Borenstein, M., Hedges, L. V., Higgins, J. P. T., & Rothstein, H. R. (2009). Introduction to meta-analysis. United Kingdom: Wiley.

*Brislin, S.J., Drislane, L.E., Smith, S.T., Edens, J.F., Patrick, C.J. (2015). Development and validation of triarchic psychopathy scales from the Multidimensional Personality Questionnaire. Psychological Assessment, 27(3), 838–851. DOI: 10.1037/pas0000087.

*Burke,J.D., Loeber, R., & Lahey, B.B. (2007). Adolescent conduct disorder and interpersonal callousness as predictors of psychopathy in young adults. Journal of Clinical Child & Adolescent Psychology, 36(3), 334–346. DOI: 10.1080/15374410701444223.

Cale, E. M., & Lilienfeld, S. O. (2002). Sex differences in psychopathy and antisocial personality disorder: A review and integration. Clinical psychology review, 22(8), 1179–1207.

Cattell, R. B., & Horn, J. L. (1978). A check on the theory of fluid and crystallized intelligence with description of new subtest designs. Journal of Educational Measurement, 15(3), 139–164.

Cleckley, H. (1941) The mask of sanity. St. Louis, MO: C.V. Mosby.

Coccaro, E. F., Berman, M. E., & Kavoussi, R. J. (1997). Assessment of life history of aggression: development and psychometric characteristics. Psychiatry research, 73(3), 147–157.

Cohen, J. (1988). Statistical power analysis for the behavioral sciences. Hillsdale, NJ: Erlbaum.

Coid, J., Yang, M., Ullrich, S., Roberts, A., & Hare, R. D. (2009). Prevalence and correlates of psychopathic traits in the household population of Great Britain. International journal of law and psychiatry, 32(2), 65–73.

Collison, K. L., Miller, J. D., & Lynam, D. R. (2019). Examining the factor structure and validity of the triarchic model of psychopathy across measures. bioRxiv. https://osf.io/7rb2v/.

Cooke, D. J., & Logan, C. (2015). Capturing clinical complexity: Towards a personality-oriented measure of psychopathy. Journal of Criminal Justice, 43(4), 262–273.

Copestake, S., Gray, N.S., & Snowden, R.J. (2011). A comparison of a self-report measure of psychopathy with the psychopathy checklist-revised in a UK sample of offenders. The Journal of Forensic Psychiatry & Psychology, 22(2), 169–182. DOI: 10.1080/14789949.2010.545134.

*Copestake, S., Gray, N.S., & Snowden, R.J. (2013). Emotional intelligence and psychopathy: A comparison of trait and ability measures. Emotion, 13(4), 691–702. DOI: 10.1037/a0031746

*de Tribolet-Hardy, F., Vohs, K., Mokros, A., & Habermeyer, E. (2014). Psychopathy, intelligence, and impulsivity in German violent offenders. International Journal of Law & Psychiatry, 37(3), 238–244.

Deary, I.J. (2013). Intelligence. Current Biology,23(16), R673–R676. DOI: 10.1016/j.cub.2013.07.021.

Debowska, A., Boduszek, D., Dhingra, K., Sherretts, N., Willmott, D., & DeLisi, M. (2018). Can we use Hare’s psychopathy model within forensic and non-forensic populations? An empirical investigation. Deviant Behavior, 39(2), 224–242.

Decuyper, M., de Pauw, S., de Fruyt, F., de Bolle, M., & de Clercq, B. (2009). A meta-analysis of psychopathy-, Antisocial PD- and FFM associations European Journal of Personality 23(7), 531–565.

*Delisi, M., Vaughn, M.G., Beaver, K.M., & Wright, J.P. (2010). The Hannibal Lecter myth: Psychopathy and verbal intelligence in the MacArthur Violence Risk Assessment Study. Journal of Psychopathology and Behavioral Assessment, 32(2), 169–177. DOI 10.1007/s10862-009-9147-z.

*Demakis, G., Rimland, C., Reeve, C., & Ward, J. (2015). Intelligence and psychopathy do not influence malingering. Applied Neuropsychology: Adult, 22(4), 262–270.

Drislane, L.E., Patrick, C.J., & Arsal, G. (2014). Clarifying the content coverage of differing psychopathy inventories through reference to the triarchic psychopathy measure. Psychological Assessment, 26(2), 350–362. DOI: 10.l37/a0035152.

Duran-Bonavila, S., Morales-Vives, F., Cosi, S., & Vigil-Colet, A. (2017). How impulsivity and intelligence are related to different forms of aggression. Personality and Individual Differences, 117, 66–70.

Duval, S.J. & Tweedie, R.L. (1998). Practical estimates of the effect of publication bias in meta-analysis. Australasian Epidemiologist, 5, 14–17.

Duval, S. & Tweedie, R. (2000) A nonparametric ‘trim and fill’ method of accounting for publication bias in meta-analysis. Journal of the American Statistical Association, 95, 89–99.

Duval, S.J. & Tweedie, R.L. (2000). Trim and fill: A simple funnel-plot-based method of testing and adjusting for publication bias in meta-analysis. Biometrics, 56, 455–463.

Edens, J. F., Marcus, D. K., Lilienfeld, S. O., & Poythress, N. G. Jr (2006). Psychopathic, not psychopath: taxometric evidence for the dimensional structure of psychopathy. Journal of abnormal psychology, 115(1), 131.

Egger, M., Smith, G. D., Schneider, M., & Minder, C. (1997). Bias in meta-analysis detected by a simple, graphical test. Bmj, 315(7109), 629–634.

*Epstein, M.K., Poythress, N.G., & Brandon, K.O. (2006). The self-report psychopathy scale and passive avoidance learning: A validation study of race and gender effects. Assessment, 13(2), 197–207. DOI: 10.1177/1073191105284992.

*Ermer, E. & Kiehl, K.A. (2010). Psychopaths are impaired in social exchange and precautionary reasoning. Psychological Science, 21(10), 1399–1405. DOI: 10.1177/0956797610384148.

*Evans, L., Ioannou, M., & Hammond, L. (2015). A predictive model of criminality in civil psychiatric populations. Journal of Criminal Psychology, 5(1), 1–12.

Farrington, D. P. (1986). Age and crime. Crime and justice, 7, 189–250.

Ferguson, C.J. (2010). Genetic contributions to antisocial personality and behavior: A meta-analytic review from an evolutionary perspective. The Journal of Social Psychology, 150(2), 160–180.

Few, L. R., Miller, J. D., Rothbaum, A. O., Meller, S., Maples, J., Terry, D. P., … & MacKillop, J. (2013). Examination of the Section III DSM-5 diagnostic system for personality disorders in an outpatient clinical sample. Journal of abnormal psychology, 122(4), 1057.

*Finn, P.R., & Hall, J. (2004). Cognitive ability and risk for alcoholism: Short-term memory capacity and intelligence moderate personality risk for alcohol problems. Journal of Abnormal Psychology, 113(4), 569–581.

*Fontaine, N., Barker, E.D., Salekin, R.T., & Viding, E. (2008). Dimensions of psychopathy and their relationships to cognitive functioning in children. Journal of Clinical Child & Adolescent Psychology, 37(3), 690–696. DOI: 10.1080/15374410802148111.

*Ford, S., Farah, M.S., Shera, D.M., & Hurt, H. (2007). Neurocognitive correlates of problem behavior in environmentally at-risk adolescents. Journal of Developmental & Behavioral Pediatrics, 28(5), 376–385. DOI: 10.1097/DBP.0b013e31811430db.

*Fowler, K.A., Lilienfeld, S.O., & Patrick, C.J. (2009). Detecting psychopathy from thin slices of behavior. Psychological Assessment, 21(1), 68–78. DOI: 10.1037/a0014938

*Frick, P.J., O’Brien, B.S., Wootton, J.M., & McBurnett, K. (1994). Psychopathy and conduct problems in children. Journal of Abnormal Psychology, 103(4), 700–707.

*Giancola, P.R., Mezzich, A.C., & Tarter, R.E. (1998). Executive cognitive functioning, temperament, and antisocial behavior in conduct-disordered adolescent females. Journal of Abnormal Psychology, 107(4), 629–641. DOI: 0021-843X/98/S3.

*Giancola, P.R., Martin, C.S., Tarter, R.E., Pelham, W.E., & Moss, H.B. (1996). Executive cognitive functioning and aggressive behavior in preadolescent boys at high risk for substance abuse/dependence. Journal of Studies on Alcohol, 57(4), 352–359.

*Gladden, P.R., Figueredo, A.J., & Jacobs,W.J. (2009). Life history strategy, psychopathic attitudes, personality, and general intelligence. Personality & Individual Differences, 46(3), 270–275. doi:10.1016/j.paid.2008.10.010.

*Goodman, R., Simonoff, E., & Stevenson, J. (1995). The impact of child IQ, parent IQ and sibling IQ on child behavioral deviance scores. Journal of Child Psychology & Psychiatry, 36(3), 409–25.

*Goodwin, E.J., Gudjonsson, G.H., Morris, R., Perkins, D., & Young, S. (2012). The relationship between sociomoral reasoning and intelligence in mentally disordered offenders. Personality & Individual Differences, 53(8), 974–979.

*Gray, J.T., Taylor, J., & Snowden, R.J. (2011). Predicting violence using structured professional judgment in patients with different mental and behavioral disorders. Psychiatry Research 187, 248–253. doi:10.1016/j.psychres.2010.10.011.

*Gray, J.T., & Snowden, R.J. (2008). Predicting violent reconvictions using the HCR–20. The British Journal of Psychiatry, 192, 384–387. DOI: 10.1192/bjp.bp.107.044065.

*Gretton, H.M., Hare, R.D., & Catchpole, R.E.H. (2004). Psychopathy and offending from adolescence to adulthood: A 10-year follow-up. Journal of Consulting and Clinical Psychology, 72(4) 636–645. DOI: 10.1037/0022-006X.72.4.636.

Gunter, T.D., Vaughn, M.G., & Philibert, R.A. (2010). Behavioral genetics in antisocial spectrum disorders and psychopathy: A review of the recent literature. Behavioral Science & The Law, 28(2), 148–173.

*Hampton, A.S., Drabick, D.A., & Steinberg, L. (2014). Does IQ moderate the relation between psychopathy and juvenile offending? Law Human Behavior, 38(1), 23–33. doi:10.1037/lhb0000036.

Hare, R. D. (1991). The Hare Psychopathy Checklist – Revised. Multi-Health Systems: Toronto, ON.

Hare, R. D. (2003). The Hare Psychopathy Checklist – Revised, 2nd edn. Multi-Health Systems: Toronto, ON.

Hare, R. D., Hart, S. D., & Harpur, T. J. (1991). Psychopathy and the DSM-IV criteria for antisocial personality disorder. Journal of abnormal psychology, 100(3), 391–398.

Hedges, L. V., & Olkin, I. (1985). Statistical methods for meta-analysis. San Diego: Academic Press.

*Heinzen, H., Köhler, D., Godt, N., Geiger, F., & Huchzermeier, C. (2011). Psychopathy, intelligence and conviction history. International Journal of Law & Psychiatry, 34(5), 336–340. doi:10.1016/j.ijlp.2011.08.002.

Hemphill, J. F. (2003). Interpreting the magnitude of correlation coefficients. American Psychologist, 58, 78–79.

Hengartner, M.P., Ajdacic-Gross,V., Rodgers, S., Müller, M., Haker, H., Rössler,W. (2014). Fluid intelligence and empathy in association with personality disorder trait-scores: Exploring the link. European Archives of Psychiatry & Clinical Neuroscience, 264(5), 441–448. DOI 10.1007/s00406-013-0441-0.

Higgins, J. P. T., Thompson, S. G., Deeks, J. J., & Altman, D. G. (2003). Measuring inconsistency in meta-analyses. British Journal of Medicine, 327, 557–560. doi.org/10.1136/bmj.327.7414.557.

*Hofvander B., Ståhlberg, O., Nydén, A., Wentz, E., degl’Innocenti, A., Billstedt, E., Forsman, A., Gillberg, C., Nilsson, T., Rastam, M.,Anckarsäter H. (2011). Life History of Aggression scores are predicted by childhood hyperactivity, conduct disorder, adult substance abuse, and low cooperativeness in adult psychiatric patients. Psychiatry Research, 30,185(1-2), 280–5. DOI: 10.1016/j.psychres.2010.05.008.

*Holland, T.R., Beckett, G.E., & Levi, M. (1981). Intelligence, personality, and criminal violence: A multivariate analysis. Journal of Consulting & Clinical Psychology, 49(1), 106–111. DOI: 0022-006X/81/4901-0106.

Isen, J. (2010). A meta-analytic assessment of Wechsler’s P>V sign in antisocial populations. Clinical Psychology Review, 30(4), 423–435.

*Jezior, K.L., McKenzie, M.E & Lee, S.S. (2016). Narcissism and callous-unemotional traits prospectively predict child conduct problems. Journal of Clinical Child & Adolescent Psychology, 45(5), 579–590. DOI: 10.1080/15374416.2014.982280.

Johnson, J. W. (2000). A heuristic method for estimating the relative weight of predictor variables in multiple regression. Multivariate Behavioral Research, 35, 1–19. doi:10.1207/S15327906MBR3501_1

Johnson, W., Carothers, A., & Deary, I. J. (2008). Sex differences in variability in general intelligence: A new look at the old question. Perspectives on Psychological Science, 3(6), 518–531.

Johnson, W., Carothers, A., & Deary, I. J. (2009). A role for the X chromosome in sex differences in variability in general intelligence?. Perspectives on Psychological Science, 4(6), 598–611.

Jones, G. (2015). Hive mind: How your nation’s IQ matters so much more than your own. Stanford: Stanford University Press.

Kavish, N., Sellbom, M., & Anderson, J. L. (2018). Implications for the Measurement of Psychopathy in the DSM–5 Using the Computerized Adaptive Test of Personality Disorder. Journal of personality assessment, 1–13. Retrieved from https://doi.org/10.1080/00223891.2018.1475393.

*Kavish, N., Bailey, C., Sharp, C., & Venta, A. (2018). On the relation between general intelligence and psychopathic traits: An examination of inpatient adolescents. Child Psychiatry Human Development, 49, 341–351.

*Kennealy, P.J., Hicks, B.M., & Patrick, C.J. (2007). Validity of factors of the psychopathy checklist-revised in female prisoners: Discriminant relations with antisocial behavior, substance abuse, and personality. Assessment, 14(4), 323–340. DOI: 10.1177/1073191107305882.

Kennedy, T. D., Burnett, K. F., & Edmonds, W. A. (2011). Intellectual, behavioral, and personality correlates of violent vs. Non-violent juvenile offenders. Aggressive Behavior, 37(4), 315–325.

*Keyes, K.M., Platt, J., Kaufman, A.S., & McLaughlin, K.A. (2017). Association of fluid intelligence and psychiatric disorders in a population-representative sample of US adolescents. JAMA Psychiatry, 74(2), 179–188. doi:10.1001/jamapsychiatry.2016.3723.

*Kipnis, D. (1965). Intelligence as a modifier of the behavior of character disorders. Journal of Applied Psychology, 49(4), 237–242.

*Klika, J.B., Herrenkohl, T.I., & Lee, J.O. (2012). School factors as moderators of the relationship between physical child abuse and pathways of antisocial behaviour. Journal of Interpersonal Violence, 28(4), 852–867. DOI: 10.1177/0886260512455865.

*Koenen, C.H., Caspi, A., Moffitt, T.E., Rijsdijk, T., & Taylor, A. (2006). Genetic influences on the overlap between low IQ and antisocial behavior in young children. Journal of Abnormal Psychology, 115(4), 787–797. DOI: 10.1037/0021-843X.115.4.787.

*Koolhof, R., Loeber, R., Wei, E.H., Pardini, D., & D’Escury, A.C. (2007). Inhibition deficits of serious delinquent boys of low intelligence. Criminal Behaviour and Mental Health, 17: 274–292. DOI: 10.1002/cbm.661.

*Kosson, D.S., Smith, S.S., & Newman, J.P. (1990). Evaluating the construct validity of psychopathy in black and white male inmates: Three preliminary studies. Journal of Abnormal Psychology, 99(3), 250–259.

Kotov, R., Krueger, R. F., Watson, D., Achenbach, T. M., Althoff, R. R., Bagby, R. M., & Zimmerman, M. (2017). The Hierarchical Taxonomy of Psychopathology (HiTOP): A dimensional alternative to traditional nosologies. Journal of Abnormal Psychology, 126(4), 454–477.

*Lahey, B.B., Loeber, R., Burke, J., & Rathouz, P.J. (2002). Adolescent outcomes of childhood conduct disorder among clinic-referred boys: Predictors of improvement. Journal of Abnormal Child Psychology, 30(4), 333–348.

LeBreton, J. M., Hargis, M. B., Griepentrog, B., Oswald, F. L., & Ployhart, R. E. (2007). A multidimensional approach for evaluating variables in organizational research and practice. Personnel Psychology, 60, 475–498. doi:10.1111/j.1744-6570.2007.00080.x

Lee, J. J., Wedow, R., Okbay, A., Kong, E., Maghzian, O., Zacher, M., … & Fontana, M. A. (2018). Gene discovery and polygenic prediction from a 1.1-million-person GWAS of educational attainment. Nature Genetics, 50(8), 1112.

Levenson, M. R., Kiehl, K. A., & Fitzpatrick, C. M. (1995). Assessing psychopathic attributes in a noninstitutionalized population. Journal of personality and social psychology, 68(1), 151–158.

Lilienfeld, S. O. (2018). The multidimensional nature of psychopathy: Five recommendations for research. Journal of Psychopathology and Behavioral Assessment, 40(1), 79–85.

Lilienfeld, S. O., Smith, S. F., Sauvigné, K. C., Patrick, C. J., Drislane, L. E., Latzman, R. D., & Krueger, R. F. (2016). Is boldness relevant to psychopathic personality? Meta-analytic relations with non-Psychopathy Checklist-based measures of psychopathy. Psychological Assessment, 28(10), 1172–1185.

Lilienfeld, S.O., & Widows, M. (2005). Professional manual for the Psychopathic Personality Inventory-Revised (PPI-R). Lutz, FL: Psychological Assessment Resources.

Lipsey, M. W., & Wilson, D. B. (2001). Practical meta-analysis. Thousand Oaks, CA: Sage.

*Loney, B.R., Frick, P.J., Clements, C.B., Ellis, M.L., & Kerlin, K. (2003). Callous-unemotional traits, impulsivity, and emotional processing in adolescents with antisocial behavior problems. Journal of Clinical Child & Adolescent Psychology, 32(1), 66–80. http://dx.doi.org/10.1207/S15374424JCCP3201_07

Lorber, M. F. (2004). Psychophysiology of aggression, psychopathy, and conduct problems: A meta-analysis. Psychological Bulletin, 130, 531–552.

*Lösel, F., & Schmucker, M. (2004). Psychopathy, risk taking, and attention: A differentiated test of the somatic marker hypothesis. Journal of Abnormal Psychology, 113(4), 522–529. DOI: 10.1037/0021-843X.113.4.522.

Lynam, D. R. (1996). Early identification of early offenders: Who is the fledgling psychopath? Psychological Bulletin, 120, 209–234.

Lynam, D. R., Hoyle, R. H., & Newman, J. P. (2006). The perils of partialling: Cautionary tales from aggression and psychopathy. Assessment, 13(3), 328–341.

*Lynam, D.R., Miller, D.J., Vachon, D., Loeber, R., Stouthamer-Loeber, M. (2009). Psychopathy in adolescence predicts official reports of offending in adulthood. Youth Violence & Juvenile Justice, 7(3), 189–207. doi:10.1177/1541204009333797.

Lynam, D., Moffitt, T., & Stouthamer-Loeber, M. (1993). Explaining the relation between IQ and delinquency: Class, race, test motivation, school failure, or self-control?. Journal of abnormal psychology, 102(2), 187–196.

*Malterer, M.B., Glass,S.J., & Newman, J.P. (2008). Psychopathy and trait emotional intelligence. Personality & Individual Differences, 44(3), 735–745.

*Masten, A.S., Hubbard, J.J., Gest, S.D., Tellegen, A., Garmezy, N., Ramirez, M., (1999). Competence in the context of adversity: Pathways to resilience and maladaptation from childhood to late adolescence. Development and Psychopathology, 11(1), 143–169.

*Mahmut, M.K., Homewood, J., & Stevenson, R.J. (2008). The characteristics of non-criminals with high psychopathy traits: Are they similar to criminal psychopaths? Journal of Research in Personality 42, 679–692.

Meldrum, R. C., Petkovsek, M. A., Boutwell, B. B., & Young, J. T. (2017). Reassessing the relationship between general intelligence and self-control in childhood. Intelligence, 60, 1–9.

Miller, J. D., & Lynam, D. R. (2012). An examination of the Psychopathic Personality Inventory’s nomological network: A meta-analytic review. *Personality Disorders: Theory, Research*, And Treatment, 3(3), 305–326.

*Moffitt, T.E., & Silva, P.A. (1988). IQ and delinquency: A direct test of the differential detection hypothesis. Journal of Abnormal Psychology, 97(3), 330–333.

*Moriconi, D.M., & Martinez, J.C. (1995). Roles of hypomania and intelligence in antisocial practices when self-esteem and family problems are considered. Psychological Reports, 76(2) 435–442.

*Morrissey, C., Hogue, T.E., Mooney, P., Lindsay, W.R., Steptoe, L., Taylor, J., & Johnston, S. (2005). Applicability, reliability and validity of the Psychopathy Checklist-Revised in offenders with intellectual disabilities: Some initial findings. International Journal of Forensic Mental Health, 4(2), 207–220.

*Nestor, P.G., Kimble, M., Berman, I., & Haycock, J. (2005). Psychosis, psychopathy, and homicide: A preliminary neuropsychological inquiry. American Journal of Psychiatry, 159(1), 138–140.

*Neumann, C.S. & Hare, R.D. (2008). Psychopathic Traits in a Large Community Sample: Links to Violence, Alcohol Use, and Intelligence. Journal of Consulting & Clinical Psychology, 76(5), 893–899. DOI: 10.1037/0022-006X.76.5.893.

*Nijman, H., Merckelbach, H., & Cima, M. (2009). Performance intelligence, sexual offending and psychopathy. Journal of Sexual Aggression, 15(3) 319–330. DOI: 10.1080/13552600903195057.

*Nomura, Y., Rajendran, K., Brooks-Gunn, J., & Newcorn, J.H. (2008). Roles of perinatal problems on adolescent antisocial behaviors among children born after 33 completed weeks: A prospective investigation. Journal of Child Psychology and Psychiatry, 49(10), 1108–1117.

*O’Boyle, E. H., Forsyth, D., Banks, G. C., Story, P. A. (2013) A meta-analytic review of the Dark Triad–intelligence connection. Journal of Research in Personality, 47(6), 789–794.

*O’Kane, A., Fawcett, D., & Blackburn, R. (1996). Psychopathy and moral reasoning: Comparison of two classifications. Personality & Individual Differences, 20(4), 505–514.

*Oscar-Berman, M., Valmas, M.M., Sawyer, K.S., Kirkley, S.M., Gansler, D.A., Merritt, D., & Couture, A. (2009). Frontal brain dysfunction in alcoholism with and without antisocial personality disorder. Neuropsychiatric Disease and Treatment, 5, 309–326.

Patrick, C. J. (2010). Operationalizing the triarchic conceptualization of psychopathy: Preliminary description of brief scales for assessment of boldness, meanness, and disinhibition. Unpublished test manual, Florida State University, Tallahassee, FL.

Paulhus, D. L., Neumann, C. S., & Hare, R. D. (2016). Manual for the Self-Report Psychopathy Scale. Toronto: Multi-Health Systems.

*Paulhus, D. L., & Williams, K. M. (2002). The dark triad of personality: Narcissism, Machiavellianism, and psychopathy. Journal of Research in Personality, 36, 556–563.

*Pera-Guardiola, V., Batalla, I., Bosque, J., Kosson, D., Pifarré, J., Hernández-Ribas, R., Goldberg, X., Contreras-Rodríguez, O., Menchón, J.M., Soriano-Mas, C., & Cardoner, N. (2016). Modulatory effects of psychopathy on Wisconsin Card Sorting Test performance in male offenders with Antisocial Personality Disorder. Psychiatry Research, 30(235), 43–48. DOI: 10.1016/j.psychres.2015.12.003.

Plomin, R., Deary, I.J. (2015). Genetics and intelligence differences: Five special findings. Molecular Psychiatry, 20(1), 98–108. DOI: 10.1038/mp.2014.105.

*Pousset, M., Tremblay, R.E., Falissard, B. (2011). Multivariate dependencies between difficult childhood, temperament and antisocial personality disorder in a population of French male prisoners. Revue d’Epidemiologie et de Sante Publique, 59(3) 169–174.

R Core Team (2013). R: A language and environment for statistical computing. R Foundation for Statistical Computing, Vienna, Austria.URL http://www.R-project.org/

*Raine, A. (1987). Effect of early environment on electrodermal and cognitive correlates of schizotypy and psychopathy in criminals. International Journal of Psychophysiology, 4(4), 277–287.

*Rispens, J., Swaab, H., Van Den Oord, E.J.C.G., Cohen-Kettenis, P., Van Engeland, H., & Van Yperen, T. (1997). WISC profiles in child psychiatric diagnosis: Sense or nonsense? Journal American Academy of Child and Adolescent Psychiatry, 36(11), 1587–1594.

Ritchie, S. J., & Tucker-Drob, E. M. (2018). How much does education improve intelligence? A meta-analysis. Psychological science, 0956797618774253.

Rogers, R., Johansen, J., Chang, J. J., & Salekin, R. T. (1997). Predictors of adolescent psychopathy: Oppositional and conduct-disordered symptoms. Journal of the American Academy of Psychiatry and the Law Online, 25(3), 261–271.

Rosenthal, R. (1979). The file drawer problem and tolerance for null results. Psychological bulletin, 86(3), 638–641.

Rosseel, Y (2012). lavaan: An R package for structural equation modeling. Journal of Statistical Software,48(2), 1–36. URL http://www.jstatsoft.org/v48/i02/

*Ruff, C.F., Templer, D.I., Ayers, J.L., Barthlow, V.L.,“WAIS Digit Span differences between prisoners and psychiatric patients. Perceptual and Motor Skills, 44,497–498.

*Salekin, R.T., Lee, Z., Schrum D., Crystal L., Kubak, F.A. (2010). Child psychopathy and protective factors: IQ and motivation to change. *Psychology*, Public Policy & Law, 16(2), 158–176.

*Salekin, R.T., Lester, W.S., Sellers, M.-K. (2004). Psychopathy in youth and intelligence: An investigation of Cleckley’s hypothesis. Journal of Clinical Child and Adolescent Psychology, 33(4), 731–742.

Salekin, R. T., Rogers, R., & Machin, D. (2001). Psychopathy in youth: Pursuing diagnostic clarity. Journal of Youth and Adolescence, 30(2), 173–195.

*Schonfeld, I.S., Shaffer, D., O’Connor, P., & Portnoy, S., (1988). Conduct Disorder and cognitive functioning: Testing three causal hypotheses. Child Development, 59 (4), 993–1007.

Schwartz, J. A., Savolainen, J., Aaltonen, M., Merikukka, M., Paananen, R., & Gissler, M. (2015). Intelligence and criminal behavior in a total birth cohort: An examination of functional form, dimensions of intelligence, and the nature of offending. Intelligence, 51, 109–118.

*Sellbom, M., & Verona, E. (2007). Neuropsychological correlates of psychopathic traits in a non-incarcerated sample. Journal of Research in Personality, 41, 276–294.

Skeem, J. L., & Mulvey, E. P. (2001). Psychopathy and community violence among civil psychiatric patients: results from the MacArthur Violence Risk Assessment Study. Journal of consulting and clinical psychology, 69(3), 358–374.

*Snowden, R., Gray, N., Smith, J., Morris, M., & MacCulloch, M. (2004): Implicit affective associations to violence in psychopathic murderers. Journal of Forensic Psychiatry & Psychology, 15(4), 620–641.

*Spironelli, C., Segrè, D., Stegagno, L., & Angrilli, A. (2013). Intelligence and psychopathy: a correlational study on insane female offenders. Psychological Medicine, 44(01), 111–116. doi:10.1017/S0033291713000615.

*Sreenivasan, S., Walker, S.C., Weinberger,L.E., Kirkish, P., & Garrick, T. (2008). Four-facet PCL–R structure and cognitive functioning among high violent criminal offenders. Journal of Personality Assessment, 90(2), 197–200. DOI: 10.1080/00223890701845476.

Sterne, J. A. & Egger, M. (2001). Funnel plots for detecting bias in meta-analysis: Guidelines on choice of axis. Journal of Clinical Epidemiology, 54, 1046–1055.

*Stevens, M.C., Kaplan, R.F., & Hesselbrock, V.M. (2003). Executive–cognitive functioning in the development of antisocial personality disorder. Addictive Behaviors, 28(2), 285–300.

Strenze, T. (2007). Intelligence and socioeconomic success: A meta-analytic review of longitudinal research. Intelligence, 35(5), 401–426.

*Strohmaier, H.N. (2015). Successful psychopathy: Do abnormal selective attention processes observed in criminal psychopaths replicate among non-criminal psychopaths? PhD Dissertation, Drexel University.

Tellegen, A. (1982). Brief manual for the Multidimensional Personality Questionnaire. Unpublished manuscript, University of Minnesota, Minneapolis.

Ttofi, M. M., Farrington, D. P., Piquero, A. R., Lösel, F., DeLisi, M., & Murray, J. (2016). Intelligence as a protective factor against offending: A meta-analytic review of prospective longitudinal studies. Journal of Criminal Justice, 45, 4–18.

*Unsworth, N., Miller, J.D., Lakey, C.E., Young, D.L., Meeks, J.T., Campbell, W.K., Goodie, A.S. (2009). Exploring the relations among executive functions, fluid intelligence, and personality. Journal of Individual Differences, 30(4), 194–200.

van Aert, R. C., Wicherts, J. M., & van Assen, M. A. (2016). Conducting meta-analyses based on p values: Reservations and recommendations for applying p-uniform and p-curve. Perspectives on Psychological Science, 11(5), 713–729.

van Aken, L., van der Heijden, P. T., van der Veld, W. M., Hermans, L., Kessels, R. P., & Egger, J. I. (2017). Representation of the Cattell–Horn–Carroll theory of cognitive abilities in the factor structure of the Dutch-Language version of the WAIS-IV. Assessment, 24(4), 458–466.

*Vaughn, M.G., DeLisi, M., Wexler, J., Barth, A., & Fletcher, J. (2011). Juvenile psychopathic personality traits are associated with poor reading achievement. Psychiatry Quarterly, 82(3), 177–190. doi:10.1007/s11126-010-9162-y.

Venales, N.C., Hall, J.R., & Patrick, C.J. (2014). Differentiating psychopathy from antisocial personality disorder: a triarchic model perspective. Psychological Medicine, 44, 1005–1013. doi:10.1017/S003329171300161X.

*Vieira, J.B., Ferreira-Santos, F., Almeida, P.R., Barbosa, F., Marques-Teixeira, J., Marsh, A.A. (2015). Psychopathic traits are associated with cortical and subcortical volume alterations in healthy individuals. Social Cognitive and Affective Neuroscience, 10(12), 1693–1704.

Viding, E., Price, T.S., Jaffee, S.R., Trzaskowski, M., Davis, O.S., Meaburn, E.L., Haworth, C.M., & Plomin, R. (2013). Genetics of callous-unemotional behavior in children. PLoS One, 8(7):e65789. DOI: 10.1371/journal.pone.0065789.

Viswesvaran, C., & Ones, D. S. (1995). Theory testing: Combining psychometric meta-analysis and structural equations modeling. Personnel Psychology, 48, 865–885. doi:10.1111/j.1744-6570.1995.tb01784.x

*Vitacco, M.J., & Kosson, D.S. (2010). Understanding psychopathy through an evaluation of interpersonal behavior: testing the factor structure of the interpersonal measure of psychopathy in a large sample of jail detainees. Psychological Assessment, 22(3), 638–649. doi:10.1037/a0019780.

*Vitacco, M.J., Neumann, C.S., & Wodushek, T. (2008). Differential relationships between the dimensions of psychopathy and intelligence. Criminal Justice & Behavior, 35(1), 48–55.

*Vitacco, M.J., Neumann, C.S., & Jackson, R.L. (2005). Testing a four-factor model of psychopathy and its association with ethnicity, gender, intelligence, and violence. Journal of Consulting & Clinical Psychology, 73(3), 466–476. DOI: 10.1037/0022-006X.73.3.466.

* Vitale, J.E., Smith, S.S., Brinkley, C.A., & Newman, J.P. (2002). The reliability and validity of the psychopathy checklist-revised in a sample of female offenders. Criminal Justice & Behavior, 29(2), 202–231.

*Wall, T.D., Sellbom, M., & Goodwin, B.E. (2013). Examination of intelligence as a compensatory factor in non-criminal psychopathy in a non-incarcerated sample. Journal of Psychopathology and Behavavioral Assessessment, 35, 450–459. DOI 10.1007/s10862-013-9358-1.

*Walters, G.D., & Kiehl, K.A. (2015). Limbic correlates of fearlessness and disinhibition in incarcerated youth: Exploring the brain behaviour relationship with the Hare Psychopathy Checklist:YouthVersion. Psychiatry Research, 230, 205–210.

*Watts, A.L., Salekin, R.T., Harrison, N., Clark, A., Waldman, I.D., Vitacco, M.J., Lilienfeld, S.O. (2016). Psychopathy: Relations with three conceptions of intelligence. Personality Disorders, 7(3),269–79. DOI: 10.1037/per0000183.

Wechsler, D. (2008). Wechsler Adult Intelligence Scale—Fourth Edition: Technical and interpretive manual. San Antonio, TX: Pearson.

*Weizmann-Henelius, G., Viemerö, V., & Eronen, M. (2004). Psychopathy in violent female offenders in Finland . Psychopathology, 37, 213–221. DOI: 10.1159/000080716.

*Welsh, G.S. (1967). Relationships of Intelligence Test Scores to Measures of Anxiety, Impulsiveness and Verbal Interests in Gifted Adolescents. Final Report. ERIC. North Carolina University: Chapel Hill.

Widiger, T. A. (2006). Psychopathy and DSM-IV psychopathology. In C. J. Patrick (Ed.), Handbook of psychopathy (pp. 156–171). New York: Guilford Press.

*Williams, K. M. (2002). Discriminating the dark triad of personality. University of British Columbia.

*Wilson, M.J., Abramowitz, C., Vasilev, G., Bozgunov, K., & Vassileva, J. (2014). Psychopathy in Bulgaria: The cross-cultural generalizability of the Hare Psychopathy Checklist. Journal of Psychopathogy & Behavioral Assessment, 36(3), 389–400. doi:10.1007/s10862-014-9405-6.

*Witt, E. (2016). The relationship of intelligence and psychopathic traits to premeditated and impulsive aggression. Eastern Washington University.

*Wodushek, T.R. (2003). Psychopathy symptom profiles and neuropsychological measures sensitive to orbitofrontal functioning. University of North Texas.

World Health Organization. (1992). International Statistical Classification of Diseases 10th Revision. Geneva: World Health Organization.

*Wright, J.P., Boisvert, B., & Vaske, J. (2009). Blood lead levels in early childhood predict adulthood psychopathy. Youth Violence and Juvenile Justice, 7(3), 208–222.

*Young, J.C., & Widom, C.S. (2014). Long-term effects of child abuse and neglect on emotion processing in adulthood. Child Abuse & Neglect, 38(8): 1369–1381. doi:10.1016/j.chiabu.2014.03.008.

*Young-Lundquist, Marcus, B.A., Boccaccini, T., & Simpler, A. (2012). Are self-report measures of adaptive functioning appropriate for those high in psychopathic traits? Behavioral Sciences and the Law, 30: 693–709. DOI: 10.1002/bsl.2039.

